# Multifaceted Functional Complexity of SARS-CoV-2 Helicase Nsp13 Underlies Its Integrated Motor and Remodeling Activities

**DOI:** 10.64898/2026.01.26.701677

**Authors:** Hai-Hong Li, Jia-Li Hou, Xue-Yang Yu, Jie Jin, Xi-Miao Hou

## Abstract

SARS-CoV-2 nonstructural protein 13 (Nsp13) is a superfamily 1 helicase essential for viral replication. Although its canonical ATP-dependent unwinding activity is well established, the broader functional repertoire of Nsp13 remains unclear. Here, we show that Nsp13 encodes a high degree of complexity by acting as a tunable nucleic acid remodeler that integrates motor and non-motor activities within a single protein. Nsp13 operates in multiple mechanistically distinct regimes, including a canonical ATP-dependent helicase mode and a Mg²⁺-primed, ATP-independent remodeling state capable of destabilizing short duplexes, hairpins, and G-quadruplexes. Mg²⁺ binding allosterically stabilizes a compact RecA1–RecA2 configuration, priming the enzyme for ATP-independent remodeling. Unwinding polarity is substrate-dependent, with duplexes supporting bidirectional remodeling, whereas G-quadruplexes are preferentially resolved in the 5′→3′ direction. Beyond strand separation, Nsp13 also exhibits robust strand annealing and nucleic acid chaperone activities. Cofactors, substrate topology, and enzyme concentration dynamically regulate these activities. Together, our findings establish Nsp13 as a highly integrated nucleic acid remodeling system and reveal how a single viral helicase switches between motor-driven and remodeling-dominated states to meet the structural demands of replication and transcription.

## Introduction

The COVID-19 pandemic, caused by Severe Acute Respiratory Syndrome Coronavirus 2 (SARS-CoV-2), has posed an unprecedented global health challenge. Although the acute phase has largely subsided, continued viral evolution and the long-term burden of post-acute sequelae (Long COVID) underscore the persistent threat posed by coronaviruses. As of December 2025, the World Health Organization has reported more than 779 million confirmed cases and over 7.1 million deaths worldwide, highlighting the ongoing need to understand the molecular mechanisms that underpin coronavirus replication and adaptability.

SARS-CoV-2 possesses large, positive-sense RNA genome (∼30 kb) that fold into highly structured and dynamic architectures[1], including stem-loops, pseudoknots[2], and G-quadruplexes (G4s)[3–5]. These RNA structures play critical roles in regulating replication, transcription, translation, and genome packaging[6]. However, the structural flexibility of RNA makes it prone to kinetic trapping in non-functional conformations[7]. This inherent complexity creates a strong dependence on nucleic acid remodeling factor that can both resolve obstructive structures and assist the formation of functional ones[8, 9]. Viral helicases and chaperones fulfill these essential functions through distinct mechanisms[10]: helicases use NTP hydrolysis to unwind structured nucleic acids, whereas chaperones promote folding or restructuring without consuming ATP.

Among the 16 nonstructural proteins encoded by SARS-CoV-2[11], Nsp13 is the only helicase and is indispensable for viral replication[12]. Nsp13 belongs to the superfamily 1b (SF1b) helicases and has been classically described as a 5′→3′ ATP-dependent motor[13, 14]. Notably, Nsp13 ranks among the most conserved coronavirus proteins[15], differing by only a single amino acid between SARS-CoV and SARS-CoV-2 (V570I), and retaining >99% identity across major viral variants[16, 17]. Such exceptional conservation suggests stringent functional constraints and implies a central, possibly multifaceted role within the viral replication–transcription complex (RTC)[18–20].

Consistent with this view, Nsp13 has traditionally been positioned at the leading edge of the RTC, where it unwinds upstream RNA, removes secondary-structure barriers, and facilitates template switching during discontinuous transcription[12, 21–23]. However, accumulating evidence indicates that its behavior cannot be fully captured by a canonical helicase model. SARS-CoV Nsp13 has been reported to exhibit ADP-stimulated chaperone-like activity that destabilizes DNA structures through transient collisions[24], while SARS-CoV-2 Nsp13 can resolve RNA stem–loops via an ATP-independent mechanism[25]. Most strikingly, recent cryo-EM analyses of the native RTC suggest that Nsp13 unwinds newly synthesized RNA duplexes with an apparent 3′→5′ polarity[26]. These findings point to a functional complexity that may exceed the conventional definition of a helicase.

Despite above intriguing findings, key questions remain unanswered. How ATP-independent remodeling by Nsp13 is activated and regulated remains unclear. Whether unwinding polarity is an intrinsic property of the enzyme or an emergent feature dictated by substrate geometry remains unclear. Moreover, its potential to recognize and remodel G4s prevalent in both host and viral genomes has not been explored.

In this study, through an integrated approach combining biochemistry, biophysics, and structural modeling, we uncover an unexpectedly high degree of functional integration within SARS-CoV-2 Nsp13. Beyond its canonical ATP-dependent helicase activity, Nsp13 engages a divalent cation–activated, ATP-independent remodeling mode that supports strand unwinding, annealing, and chaperone-like restructuring across diverse nucleic acid substrates, including duplexes, hairpins, and G4s. These activities are dynamically tuned by cofactor availability, substrate topology, and enzyme concentration, enabling Nsp13 to switch between motor-driven and remodeling-dominated states. By establishing how multiple, mechanistically distinct nucleic acid transactions are integrated within a single viral helicase, our work expands the conceptual framework for helicase function and highlights new principles for understanding—and potentially targeting—viral replication machineries.

## Results

### Nsp13 is a broad-spectrum nucleic acid-binding protein

SARS-CoV-2 Nsp13 adopts a conserved helicase architecture, comprising an N-terminal zinc-binding domain (ZBD), two RecA-like domains (1A and 2A), a bridging 1B domain, and an auxiliary stalk domain that connects the ZBD to the helicase core (**Figure 1A**; PDB: 7nio)[28]. The ATP-binding pocket lies at the cleft between domains 1A and 2A, whereas the nucleic-acid–binding groove spans domains 1A, 2A, and 1B. To biochemically characterize the protein, we expressed and purified recombinant SARS-CoV-2 Nsp13 (hereafter referred to as Nsp13) in *E. coli* (**Figure 1B**). LC–MS/MS analysis confirmed protein identity, achieving 87% sequence coverage with >93% of peptide–spectrum matches uniquely assigned to SARS-CoV-2 Nsp13 (**Table S2**). Size-exclusion chromatography further indicated that Nsp13 elutes as a single symmetric peak consistent with a monomeric state in solution (**Figure 1C**).

**Figure 1.**
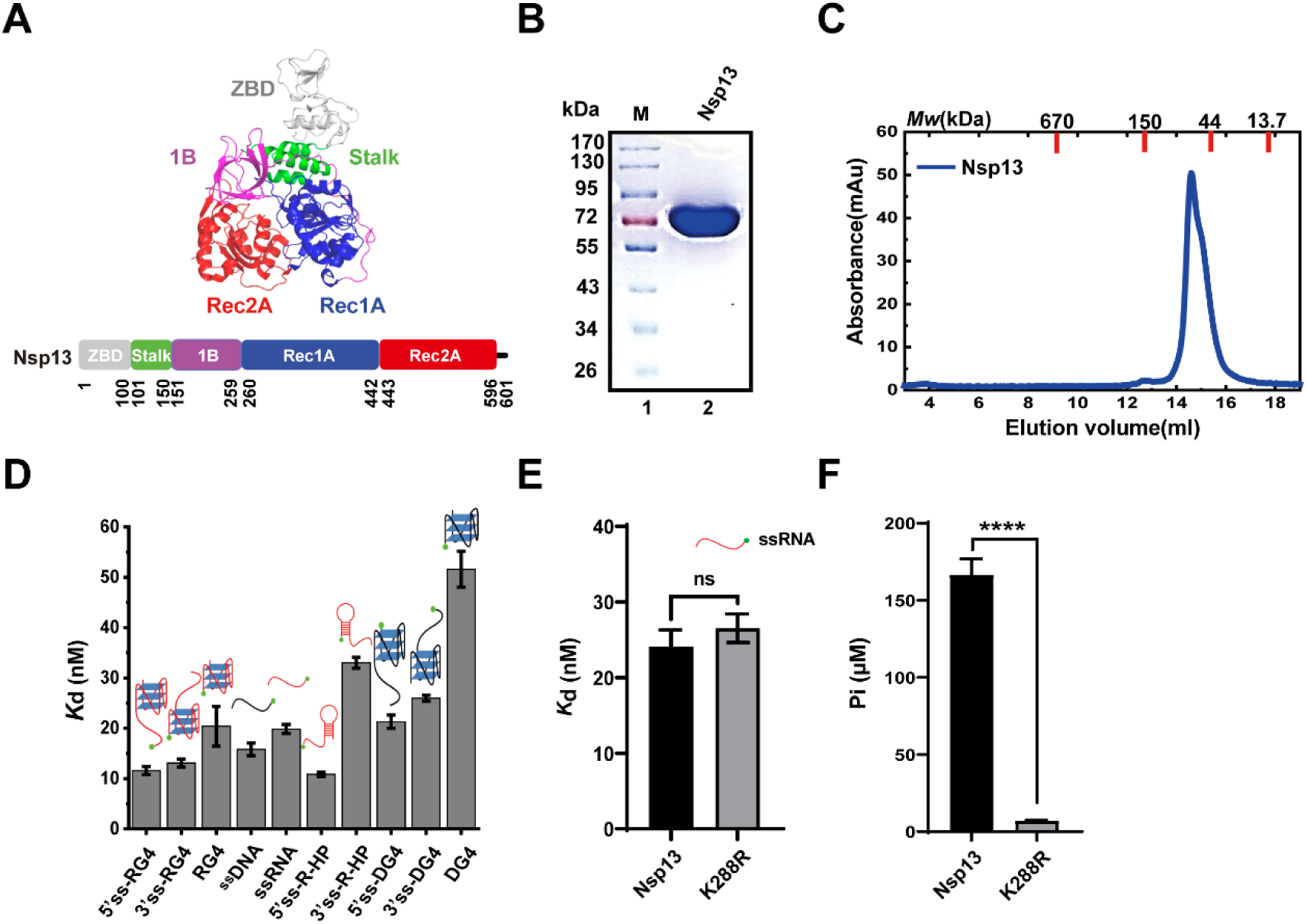
Structural features and biochemical characterization of SARS-CoV-2 Nsp13. (A) Domain architecture and structure of Nsp13 from PDB (7nio). (B) SDS-PAGE analysis of purified Nsp13. (C) Size-exclusion chromatography (SEC) profile of Nsp13, indicating a monomeric state in solution. (D) Equilibrium dissociation constants (Kd) of Nsp13 with various nucleic acid substrates. (E) Binding affinity of the ATPase-deficient mutant K288R compared to wild-type Nsp13. (F) ATPase activities of wild-type Nsp13 and K288R mutant in the presence of ssRNA. Data are presented as mean ± SD (n=3). Statistical significance was determined by unpaired Student’s t-test for comparisons between two groups, or one-way ANOVA followed by Tukey’s HSD test for comparisons involving more than two groups; ns, P>0.05, P < 0.05, p < 0.01, P<0.001, P<0.0001 (apply to all figures).

Fluorescence anisotropy assays showed that Nsp13 binds a broad spectrum of DNA and RNA substrates with low- to mid-nanomolar affinities (**Figure 1D; Table S3**). ATP hydrolysis was not required for substrate engagement, as the ATPase-deficient K288R mutant [29] displayed binding affinities indistinguishable from the wild-type protein (**Figure 1E–F**). Specifically, Nsp13 bound single-stranded DNA (Kd = 15.8 ± 1.3 nM) and RNA (Kd = 19.9 ± 0.9 nM) with comparable affinity, but preferentially recognized structured RNA elements, including a 5′-tailed RNA hairpin (Kd = 10.9 ± 0.4 nM) and an RNA G4 (Kd = 11.6 ± 0.8 nM). Notably, most binding curves were better fitted by a Hill model with coefficients >1 (**Figure S1**), indicating cooperative binding behavior[27].

Collectively, these findings establish SARS-CoV-2 Nsp13 as a potent, broad-specificity nucleic acid–binding helicase, with a marked preference for structured RNA elements.

### Divalent cations activate a novel ATP-independent DNA unwinding mode

To comprehensively assess the unwinding capability of Nsp13, we initially employed a 16-bp duplex DNA substrate with a 14-nucleotide 5’overhang (5’Oh_S14D16_), consistent with its reported 5’→3’ polarity[30]. Surprisingly, in the absence of ATP, Nsp13 efficiently unwound this substrate in a concentration-dependent manner when Mg²⁺ was present (5 mM; **Figure 2A**). This activity exhibited a clear dependence on Mg²⁺ concentration over a broad range (0–20 mM; **Figure 2B**) and was retained in an N-terminal His-tag–deleted construct, excluding tag-dependent artifacts (**Figure S2**).

**Figure 2.**
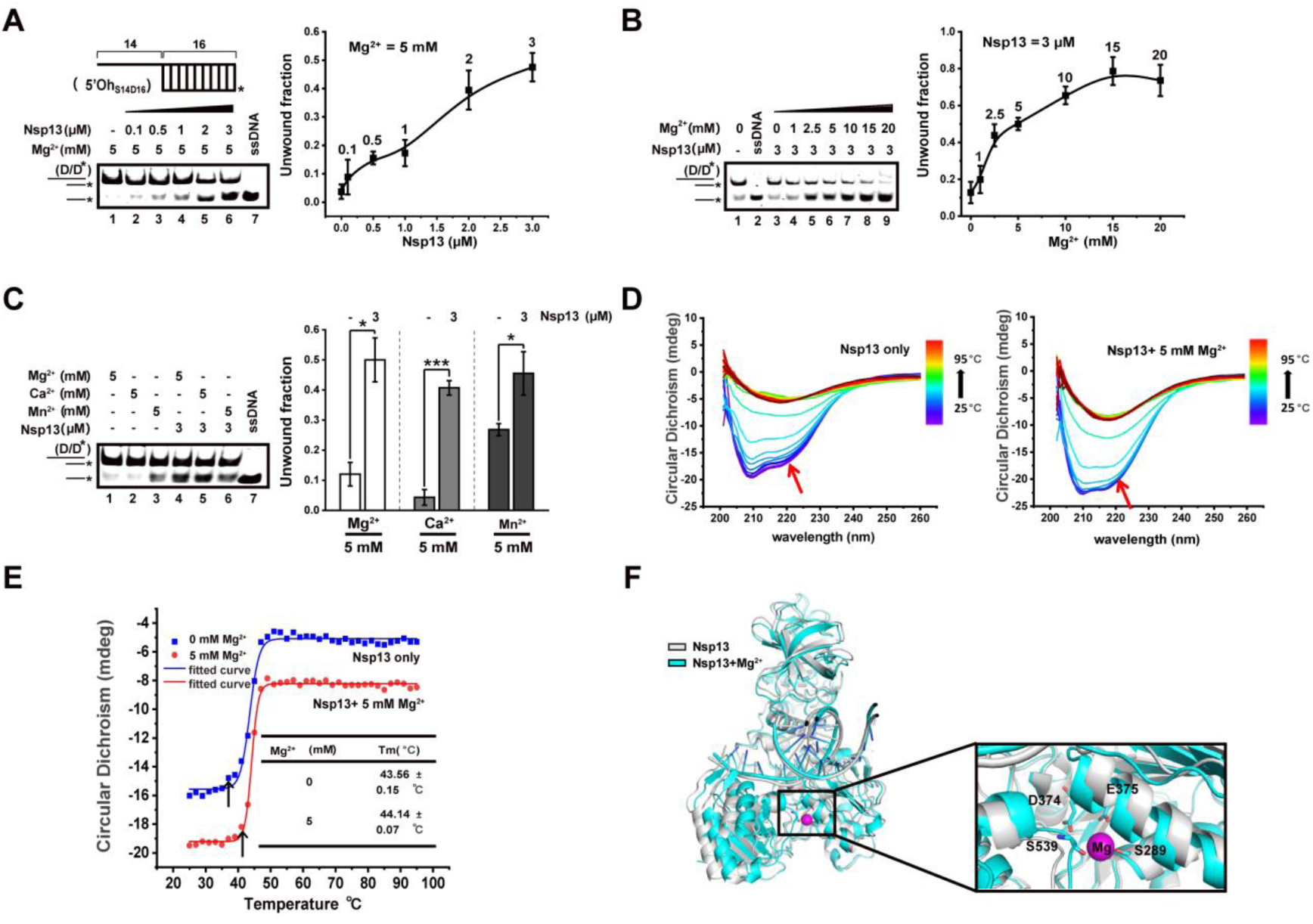
Divalent cations activate ATP-independent DNA unwinding and allosterically stabilize Nsp13. (A) Schematic of the 5′-overhang DNA substrate with **^*^** denoting the FAM label (5′Oh_S14D16_) and EMSA showing Nsp13-mediated, concentration-dependent unwinding in the presence of 5 mM Mg²⁺ without ATP (DNA substrate: 40 nM). Right panel: quantification of unwound DNA fraction. (B) EMSA and quantification illustrating the effect of Mg²⁺ concentration (0–20 mM) on ATP-independent DNA unwinding. (C) EMSA and quantification comparing the activation of DNA unwinding by different divalent cations (Mg²⁺, Ca²⁺, Mn²⁺). (D) CD spectra of Nsp13 in the absence and presence of 5 mM Mg²⁺. (E) Thermal denaturation curves of Nsp13 monitored by CD at 222 nm. (F) AlphaFold3-predicted structural models of the Nsp13–RNA fork complex, highlighting Mg²⁺-induced compaction between the RecA1 and RecA2 domains (RNA was selected for modeling due to its higher prediction confidence relative to DNA). Data are presented as mean ± SD (n=3).

This ATP-independent activity was not restricted to Mg²⁺. Other divalent cations, including Ca²⁺ and Mn²⁺, also activated Nsp13 with a relative efficacy of Mg²⁺ > Ca²⁺ > Mn²⁺ (**Figure 2C**). Together, these results reveal a previously unrecognized DNA-remodeling activity of Nsp13 that is strongly activated by divalent cations and does not require ATP. This cation-driven unwinding mode represents a mechanistically distinct operational state of the enzyme, markedly different from its canonical ATP-fueled helicase cycle.

### Mg²⁺ allosterically stabilizes Nsp13 and primes a non-processive remodeling conformation

To elucidate the mechanism underlying Mg²⁺-activated ATP-independent unwinding, we first examined whether Mg²⁺ directly destabilizes the duplex substrate. FRET melting assays revealed that 5 mM Mg²⁺ significantly stabilized the double-stranded DNA, increasing its melting temperature (Tₘ) from 49 °C to 57.7 °C (**Figure S3A**). Addition of Nsp13 up to 1.8 µM only induced a marginal decrease in Tₘ (55.5-56.4°C; **Figure S3B-C**), indicating that the observed unwinding is not attributable to substrate destabilization. Moreover, fluorescence anisotropy measurements showed that Mg²⁺ modestly weakened Nsp13’s binding affinity for a 5′-overhang DNA substrate, with the Kd increasing from 265.5 ± 0.41 nM (0 mM Mg²⁺) to 434.4 ± 4.23 nM (0.5 mM Mg²⁺) (**Figure S3D**), ruling out enhanced binding as the driver of unwinding.

We next asked whether Mg²⁺ directly alters the structural stability and conformational state of Nsp13. Circular dichroism (CD) thermal unfolding experiments revealed that Mg²⁺ modestly stabilized Nsp13 at near-physiological temperatures, despite no significant change in overall thermostability (**Figure 2D-E**). In the absence of Mg²⁺, unfolding began at approximately 37 °C, whereas the onset of unfolding shifted to ∼41 °C in the presence of Mg²⁺, suggesting that Mg²⁺ helps maintain a more compact conformation under near-physiological conditions.

AlphaFold3 structural modeling provided further mechanistic insight. As shown in **Figure S4A**, the predicted Nsp13 structure closely matched the known crystal structure, supporting the validity of our modeling. Notably, Mg²⁺ binding promotes a more compact arrangement between the RecA1 and RecA2 domains, primarily through interactions involving several key amino acids (**Figure 2F**), offering a structural explanation for the thermal stabilization observed in CD melting. This model of cation-driven allosteric stabilization is further supported by the conformational behavior of Nsp13 in the presence of ADP-Mg²⁺, where the protein adopted a more compact state with specific residues potentially involved in Mg²⁺ coordination (**Figure S4B**).

Together, these findings establish Mg²⁺ as an allosteric effector that stabilizes Nsp13 and promotes a compact, low-processivity conformational state. This Mg²⁺-primed state provides a mechanistic basis for the ATP-independent unwinding mode and highlights how Nsp13 can toggle between distinct functional regimes depending on the availability of nucleotides and cations.

### Energy availability governs switching between distinct DNA remodeling modes and polarities

We next investigated the role of nucleotide binding and hydrolysis. We found that the non-hydrolyzable ATP analog AMP-PNP could stimulate dsDNA unwinding similarly as ATP and more effectively than Mg²⁺ alone (**Figure 3A-B**). Remarkably, K288R exhibited even greater unwinding activity in the presence of ATP than wild-type Nsp13 under either ATP or AMP-PNP conditions. AlphaFold3 modeling suggests that this enhancement may reflect a more tightly closed RecA1–RecA2 conformation in the ATP–Mg²⁺–bound K288R mutant (**Figure S5**). These findings demonstrate that ATP binding, even without hydrolysis, is sufficient to induce a conformational state in Nsp13 that is highly proficient in DNA unwinding.

**Figure 3.**
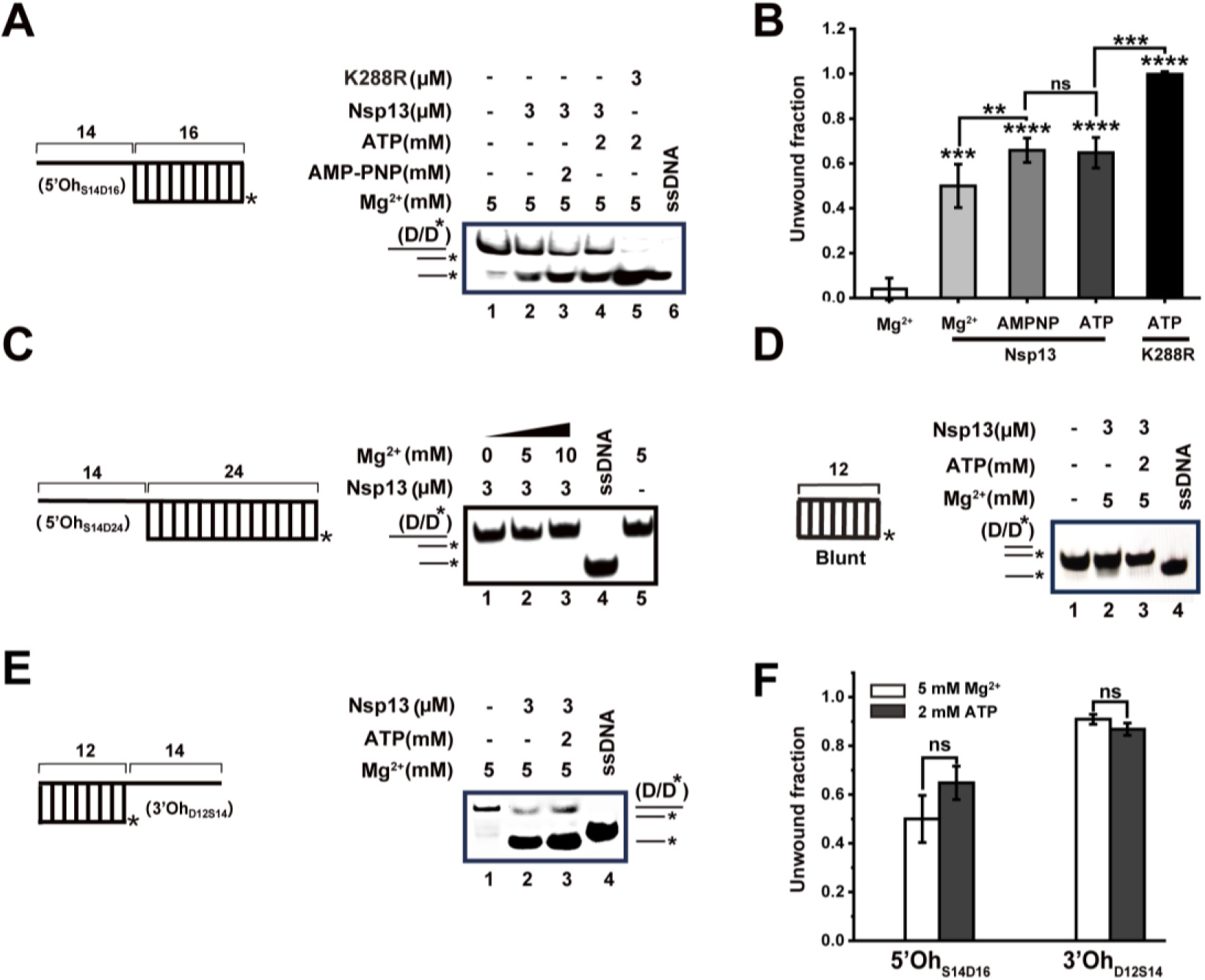
Energy- and substrate-dependent DNA unwinding by Nsp13. (A–B) EMSA and quantification comparing Nsp13-mediated duplex DNA unwinding under different nucleotide and Mg²⁺ conditions. K288R was included as a control to assess the contribution of ATP hydrolysis. (C) Schematic of the 24-bp DNA substrate unwound only in the presence of Mg^2+^. (D) Blunt-ended duplex DNA cannot be unwound by Nsp13 under either ATP-independent or ATP-dependent conditions. (E) Schematic of a 3′-overhang DNA substrate (3′Oh_D12S14_) and EMSA showing Nsp13-mediated unwinding in the presence of 5 mM Mg²⁺, either in the absence or presence of 2 mM ATP. (F) Quantification of unwound DNA fraction. Data represent mean ± SD (n = 3).

The ATP-independent unwinding mode, however, has limited processivity. When challenged with a more stable 24-bp dsDNA substrate, Nsp13 failed to unwind it in the presence of Mg²⁺ alone (**Figure 3C**), requiring ATP hydrolysis for this more demanding task. This indicates that while metal ion activation enables basal remodeling activity, processive unwinding of stable substrates depends on energy derived from ATP hydrolysis. Furthermore, the ATP-dependent activity was sensitively regulated by the ATP/Mg²⁺ balance. At a fixed ATP concentration (1 mM), excess Mg²⁺ beyond 1 mM strongly inhibited unwinding (**Figure S6A**). Conversely, maintaining a 1:1 ATP/Mg²⁺ ratio preserved robust unwinding efficiency across a range of ATP concentrations (1-5 mM; **Figure S6B**), underscoring the critical balance between these cofactors for optimal Nsp13 function.

Having defined two distinct unwinding regimes—an ATP-independent, cation-activated mode with limited processivity and a canonical ATP hydrolysis–dependent mode capable of processive strand separation—we next asked whether these modes impose a fixed directionality on DNA unwinding. Nsp13 has traditionally been classified as a strict 5′→3′ helicase based primarily on assays employing substrates with 5′ single-stranded overhangs. To test whether this polarity represents an intrinsic constraint or a context-dependent property, we examined Nsp13 activity on duplex DNA substrates bearing a 3′ single-stranded overhang.

As a boundary condition, blunt-ended duplex DNA was not unwound under either ATP-independent or ATP-dependent conditions, confirming the requirement for a single-stranded loading region (**Figure 3D**). In contrast, when a substrate containing a 12-bp duplex region and a 3′ overhang was used, we observed robust unwinding in the presence of Mg²⁺ alone, indicating that the cation-activated, ATP-independent mode is sufficient to support efficient 3′→5′ DNA unwinding (**Figure 3E-F**). Addition of ATP produced a comparable extent of unwinding, suggesting functional redundancy between the ATP-independent and ATP-dependent modes on this substrate. Similar results were obtained with a more stable substrate containing a 16-bp duplex region (**Figure S6C**).

Together, these results establish that DNA unwinding by Nsp13 is governed by two orthogonal parameters: the energetic state of the enzyme and the geometry of the DNA substrate. Mg²⁺ binding induces a compact conformation that primes Nsp13 for basal strand remodeling but supports only limited processivity. In contrast, ATP binding alone is sufficient to promote a highly active unwinding conformation, while ATP hydrolysis provides the energy required to sustain processive separation of more stable duplexes. Importantly, neither the cation-activated nor the ATP-dependent mode enforces a strict polarity constraint. Instead, unwinding directionality emerges from the substrate context, revealing pronounced mechanistic plasticity that enables Nsp13 to unwind duplex DNA in either direction. This functional flexibility provides an essential framework for interpreting recent structural studies showing that, within the native RTC, Nsp13 operates with 3′→5′ polarity to unwind newly synthesized RNA duplexes[26].

### RNA substrates reveal a Mg²⁺- and concentration-dependent remodeling regime distinct from DNA

Having established the ATP-independent unwinding activity of Nsp13 on DNA, we sought to determine if a similar mechanism applies to RNA substrates.

A key finding was the narrow optimal range for Mg²⁺ in facilitating RNA unwinding. When assayed using an RNA-fork substrate (**Figure 4A, upper panel**), Nsp13 (2 µM) exhibited negligible unwinding activity in the absence of Mg²⁺, and the substrate showed signs of instability under Mg²⁺-free condition (**Figure S7A**). Strikingly, the introduction of 0.5 mM Mg^2+^ enabled a pronounced unwinding activity, with efficiency peaking at ∼30% (**Figure 4A-B**). K288R exhibited a comparable level of activity at this Mg²⁺ concentration (**Figure 4C-D**, lane 5), confirming that this pathway does not require ATP hydrolysis. However, this activation was highly specific, as increasing the Mg²⁺ concentration to 1 mM markedly suppressed the activity to 15.4% (**Figure S7A**, **Figure 4B**). This sharp decline stands in stark contrast to the positive correlation between Mg²⁺ concentration and unwinding efficiency observed for duplex DNA (**Figure 2B**), highlighting a nucleic acid-type-specific regulatory mechanism.

**Figure 4.**
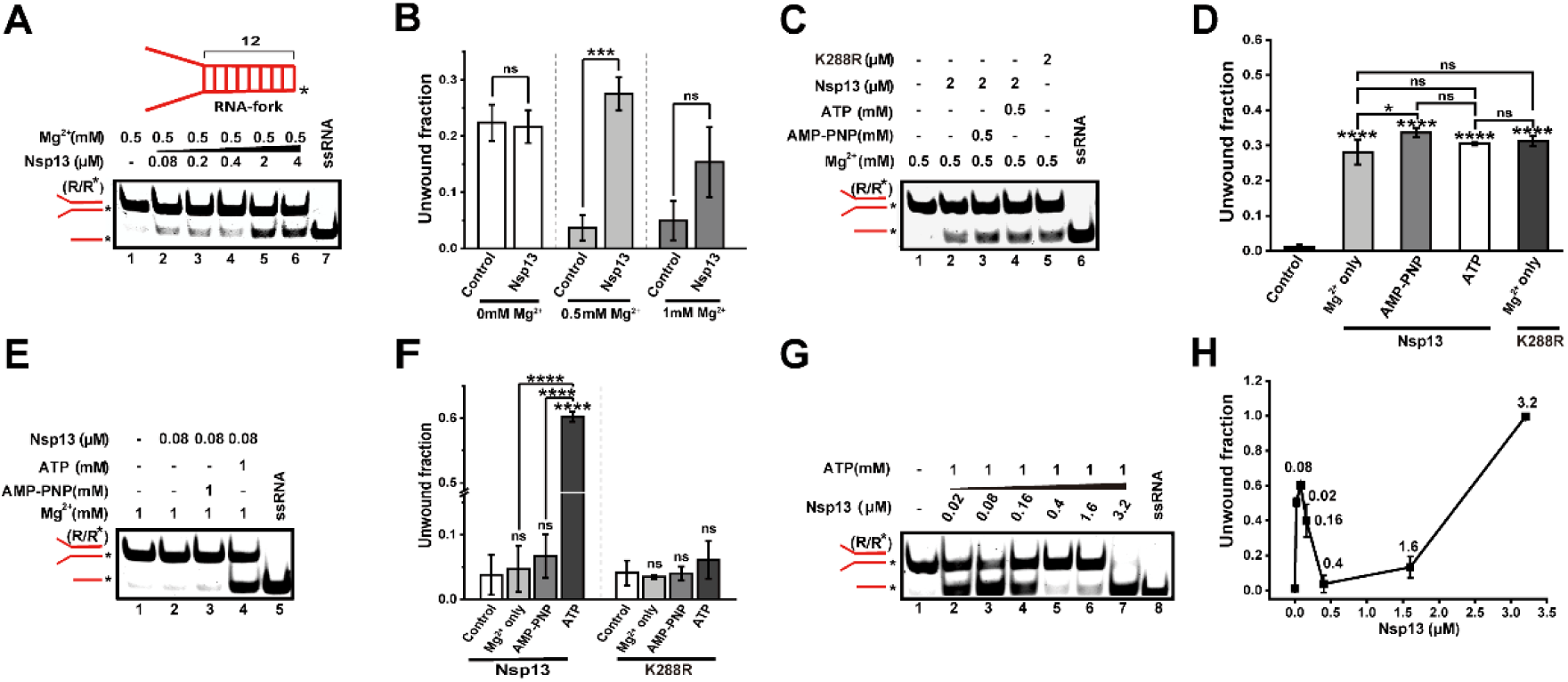
Mg²⁺ activates ATP-independent and -dependent RNA unwinding with unique sensitivity and concentration dependence. (A) Schematic of the RNA-fork substrate and EMSA showing RNA unwinding at 0.5 mM Mg²⁺ without ATP. (B) Quantification of unwinding efficiency by 2 µM Nsp13 at 0, 0.5, and 1 mM Mg²⁺. (C-D) EMSA and quantification of RNA unwinding under different ATP conditions. (E-F) Comparison of wild-type Nsp13 and K288R mutant unwinding at low protein concentration (80 nM). (G-H) EMSA and quantification showing triphasic concentration dependence of ATP-dependent RNA unwinding at 1 mM Mg²⁺. RNA substrate concentration: 40 nM. Data are presented as mean ± SD (n=3).

The interplay between nucleotide binding and the cation activation also differed for RNA. While AMP-PNP stimulated dsDNA unwinding, it did not enhance dsRNA unwinding beyond the level achieved by 0.5 mM Mg²⁺ alone (**Figure 4C-D**). Intriguingly, at 1 mM Mg^2+^ where ATP-independent mode is insignificant, at a low protein concentration (80 nM), a striking difference emerged: ATP-dependent unwinding (60%) overwhelmingly dominated, showing a ∼13-fold enhancement over the minimal activity achieved with Mg²⁺ only (4.7%) (**Figure 4E-F**). The ATP hydrolysis-deficient mutant K288R showed no differential activity across treatments (**Figure S7B, Figure 4F**), confirming that it is ATP hydrolysis that serves as the key and potent trigger for this highly efficient RNA unwinding mode at low enzyme concentrations.

We further characterized the concentration dependence of ATP-dependent RNA unwinding at 1 mM Mg²⁺ (**Figure 4G**). Strikingly, it shows triphasic pattern-activation, inhibition, and restoration (**Figure 4H**), consistent with that was observed for the ATP-independent pathway (**Figure S7C**). At low Nsp13 concentrations (≤ 160 nM), robust RNA unwinding was observed. However, at an intermediate concentration of 0.4 µM, the unwinding activity was significantly inhibited. Further increasing the Nsp13 concentration beyond 1.6 µM led to a pronounced recovery of unwinding efficiency (**Figure 4H**). Control experiments confirmed that this inhibitory phase is governed by the absolute concentration of Nsp13 rather than the protein-to-substrate ratio (**Figure S8**), suggesting an intrinsic property of Nsp13 at this specific concentration range. This nonlinear behavior was specific to RNA substrates, as DNA unwinding exhibited a standard, cooperative increase with protein concentration under identical conditions (**Figure S9**).

Altogether, Nsp13 employs a distinct and sophisticated strategy to unwind RNA, characterized by a narrow Mg²⁺ optimum and a complex concentration-dependent regulatory mechanism. This behavior, which is qualitatively different from its action on DNA, suggests that RNA substrates induce specific conformational changes or functional states in Nsp13, leading to a unique regulatory mechanism tailored for RNA processing.

### Nsp13 resolves G4 structures through dual ATP-dependent and -independent pathways

Having established that Nsp13 exhibits versatile nucleic acid remodeling activities on conventional duplexes, we next sought to investigate its capacity to resolve highly stable G4 structures. Given the prevalence of putative G4-forming sequences in both the human and SARS-CoV-2 genomes and their potential regulatory roles in viral replication[31, 32], we evaluated Nsp13’s activity on human telomeric DNA G4 (DG4) [33]and a conserved viral RNA G4 (RG-1) [6]using a gel-based unfolding assay (**Figure 5A**).

**Figure 5.**
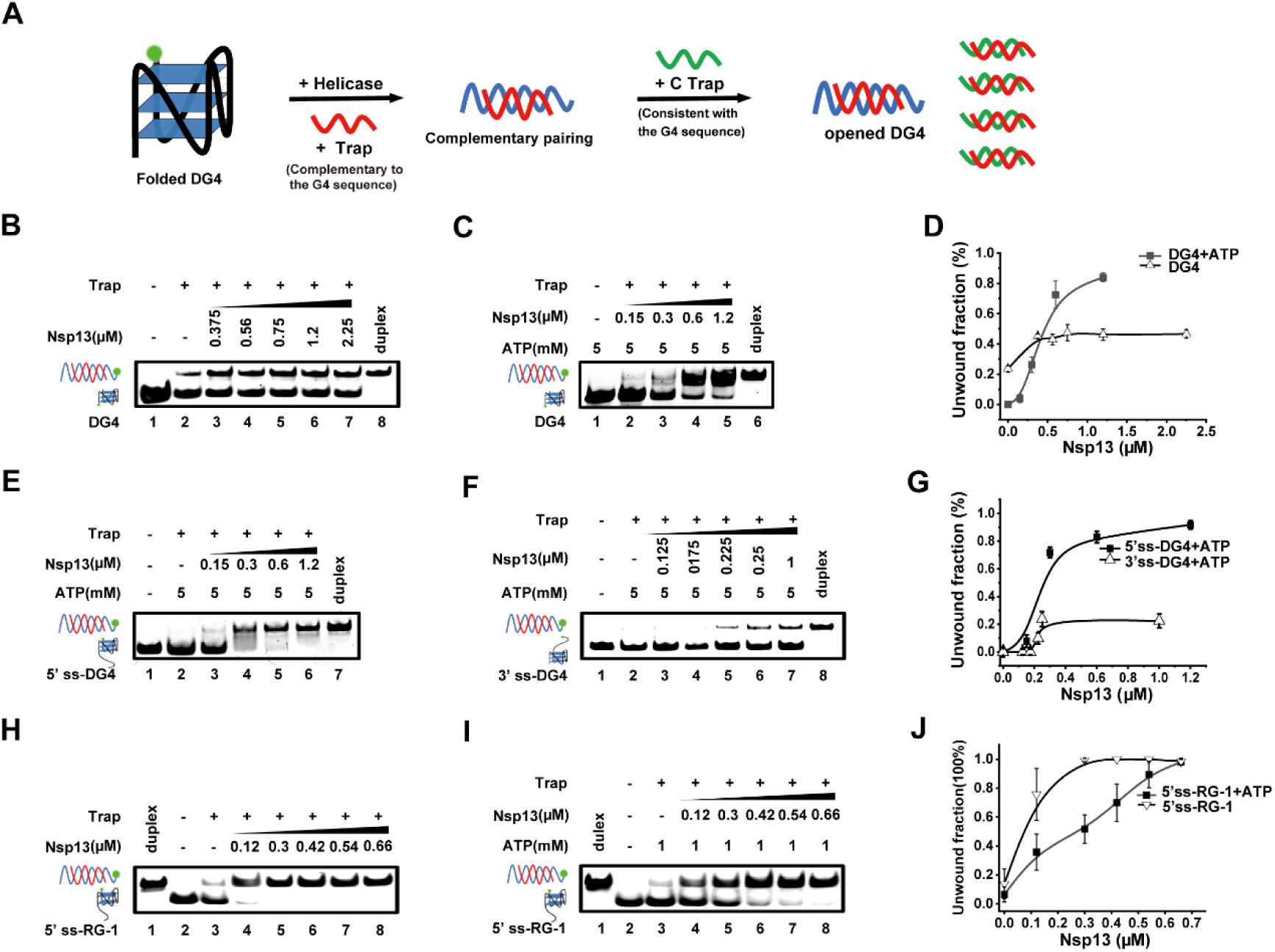
Nsp13 unwinds G4 structures through dual ATP-dependent and -independent pathways. (A) Schematic of the G4 unfolding assay. (B–C) EMSA showing ATP-independent and ATP-dependent unfolding of blunt-ended DG4 (30 nM). (D) Quantification of DG4 unfolding efficiency. (E–F) EMSA showing direction-dependent unwinding of 5′- and 3′-overhang DG4 (25 nM) substrates. (G) Quantification confirming 5′→3′ directionality. (H–I) EMSA showing ATP-independent and ATP-dependent unfolding of SARS-CoV-2 RNA G4 (RG-1, 60 nM). (J) Quantification of RG-1 unfolding efficiency.

Strikingly, Nsp13 unwound the blunt-ended DG4 structure even in the absence of ATP, with efficiency increasing in a concentration-dependent manner (**Figure 5B**). The addition of ATP further enhanced the unfolding proportion (**Figure 5C, D**), demonstrating that Nsp13 possesses dual ATP-independent and ATP-dependent G4 resolving activities. **Figure 5E–G** further showed that Nsp13 efficiently unwound the 5′-overhang DG4 in an ATP-dependent manner, but showed much weaker activity on the 3′-overhang substrate. These results indicate that, although Nsp13 can switch polarity in duplex DNA unwinding, its G4 unfolding activity predominantly follows a 5′→3′ direction.

This dual-mode capability also extended to RNA G4 structures. Nsp13 efficiently unfolded the SARS-CoV-2 genomic RG-1 through both ATP-independent and ATP-dependent mechanisms (**Figure 5H-J**). Intriguingly, the ATP-independent activity surpassed the ATP-dependent one across a broad protein concentration range (**Figure 5J**). This preference likely reflects intrinsic stability differences: RG-1 forms a relatively less stable two-layer G4 [34]that is more easily disrupted by direct binding–induced destabilization (chaperone mode), whereas the more stable three-layer DG4 requires processive, ATP-driven unwinding. Consistent with duplex RNA results (**Figure 4G-H**), RG-1 unfolding was inhibited at high Nsp13 concentrations (**Figure S10**), indicating a shared concentration-dependent regulatory behavior.

These findings significantly expand the functional repertoire of Nsp13, positioning it as a dual-mode nucleic acid remodeler capable of resolving complex G4 structures prevalent in both host and viral genomes. This activity, fine-tuned by ATP availability and substrate architecture, likely plays a crucial role in managing genomic stability and facilitating the progression of the viral RTC through G-rich regions.

### Nsp13 functions as a unified ATP-independent nucleic acid remodeler across diverse substrates

Based on the comprehensive characterization of Nsp13’s ATP-independent unwinding activities on duplex DNA, RNA, and G4 (**Figures 2-5**), we further investigated whether it possesses broader nucleic acid remodeling functions[9].

Nsp13 demonstrated potent strand-annealing activity. It efficiently facilitated the hybridization of complementary single-stranded DNA into various duplex structures-including those with 5’-overhang, 3’-overhang, and fork-without any requirement for ATP (**Figure 6A-C; Figure S11A-D**). Systematic analysis across multiple DNA substrates revealed a consistent biphasic dependence of annealing efficiency on Nsp13 concentration, particularly evident in the presence of Mg²⁺ alone. The activity increased sharply at low nanomolar concentrations, reached an optimum at approximately 200 nM, and then declined significantly with further increases in protein concentration (**Figure 6B, S11A-D**). This non-monotonic pattern suggests a self-regulatory mechanism, potentially arising from non-productive substrate sequestration or protein oligomerization at elevated concentrations. Crucially, the simultaneous presence of ATP and Mg²⁺ switched Nsp13’s function from annealing to rapid unwinding, highlighting its dynamic operational mode (**Figure 6B-C**).

**Figure 6.**
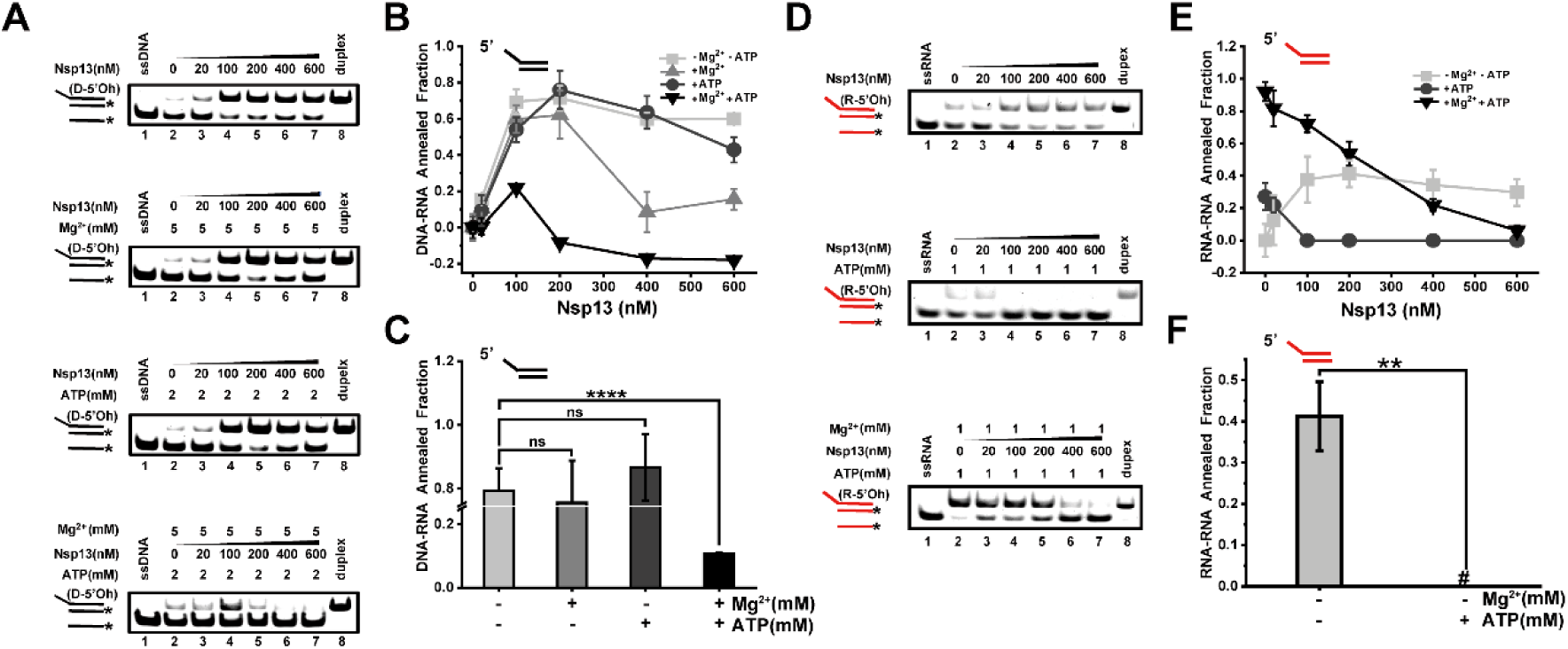
Nsp13 exhibits ATP/Mg²⁺-independent nucleic acid annealing activity. (A) EMSA of DNA strand annealing by Nsp13 under four cofactor conditions: no Mg²⁺/ATP, Mg²⁺ only, ATP only, and Mg²⁺+ATP. (B) Concentration-response curve showing biphasic DNA annealing. (C) Quantification of DNA annealing efficiency at 200 nM Nsp13. (D) EMSA of RNA strand annealing under three cofactor conditions: no Mg²⁺/ATP, ATP only, and Mg²⁺+ATP. The Mg²⁺-only condition was omitted because 1 mM Mg²⁺ promotes spontaneous hybridization of the single-stranded RNA substrates. (E) Concentration-response curve of RNA annealing. (F) Quantification of RNA annealing efficiency at 200 nM Nsp13. Data are presented as mean ± SD (n=3).

In parallel, Nsp13 exhibited RNA strand-annealing activity with distinct substrate specificity and cofactor sensitivity. In the absence of both Mg²⁺ and ATP, Nsp13 efficiently promoted annealing of RNA substrates with 5’-overhang and fork structures, while showing minimal activity on 3’-overhang or blunt ends (**Figure S11E-G**). The introduction of 1 mM ATP alone significantly suppressed this annealing activity (**Figure 6D-F**). Notably, in the presence of 1 mM Mg²⁺, where substantial spontaneous RNA duplex formation occurred, the addition of 1 mM ATP triggered a functional switch from annealing to robust unwinding (**Figure 6D-E**). This cofactor-dependent transition reveals a finely tuned dynamic equilibrium that is uniquely adapted for RNA substrate remodeling.

To further examine its chaperone function, we employed a classic strand-exchange assay[35] (**Figure S12A**). When incubated with Nsp13 in the absence of ATP, pre-folded DNA and RNA hairpins were efficiently destabilized, leading to a time-dependent formation of a more stable, hybridized duplex (**Figure S12B-C**). This remodeling was more efficient for DNA than RNA, consistent with the annealing results. The failure of a control 5′→3′ helicase, ToPif1[36], to catalyze this reaction under identical conditions underscored the uniqueness of Nsp13’s inherent chaperone activity (**Figure S12D**).

In conclusion, our work identifies a previously unreported fundamental aspect of SARS-CoV-2 Nsp13: a unified, ATP-independent nucleic acid remodeling function encompassing both strand annealing and secondary structure destabilization. This activity, operating alongside canonical ATP-dependent helicase function and modulated by cations, protein concentration, and substrate identity, provides the viral replication machinery with exceptional flexibility to manage complex nucleic acid structures throughout the viral life cycle.

## Discussion

In this study, we redefine SARS-CoV-2 Nsp13 as a highly integrated, mode-switchable nucleic acid remodeler whose activity is dynamically regulated by cofactors, substrate architecture, and enzyme concentration. Rather than functioning solely as a canonical ATP-dependent helicase, Nsp13 coordinates motor-driven and ATP-independent remodeling activities within a single protein to manage structurally complex nucleic acids during viral replication and transcription.

Our results establish Nsp13 as an extreme example of functional integration within the helicase family, challenging the traditional view that helicase function is defined primarily by ATP-driven, unidirectional unwinding. In addition to its canonical helicase mode, Nsp13 destabilizes duplexes, hairpins, and G4s through a divalent cation–activated, ATP-independent pathway and catalyzes strand annealing and chaperone-like restructuring. These activities are not independent but are coordinated and selectively engaged by physicochemical cues such as Mg²⁺ availability, substrate topology, and protein concentration (**Figure 7**).

**Figure 7.**
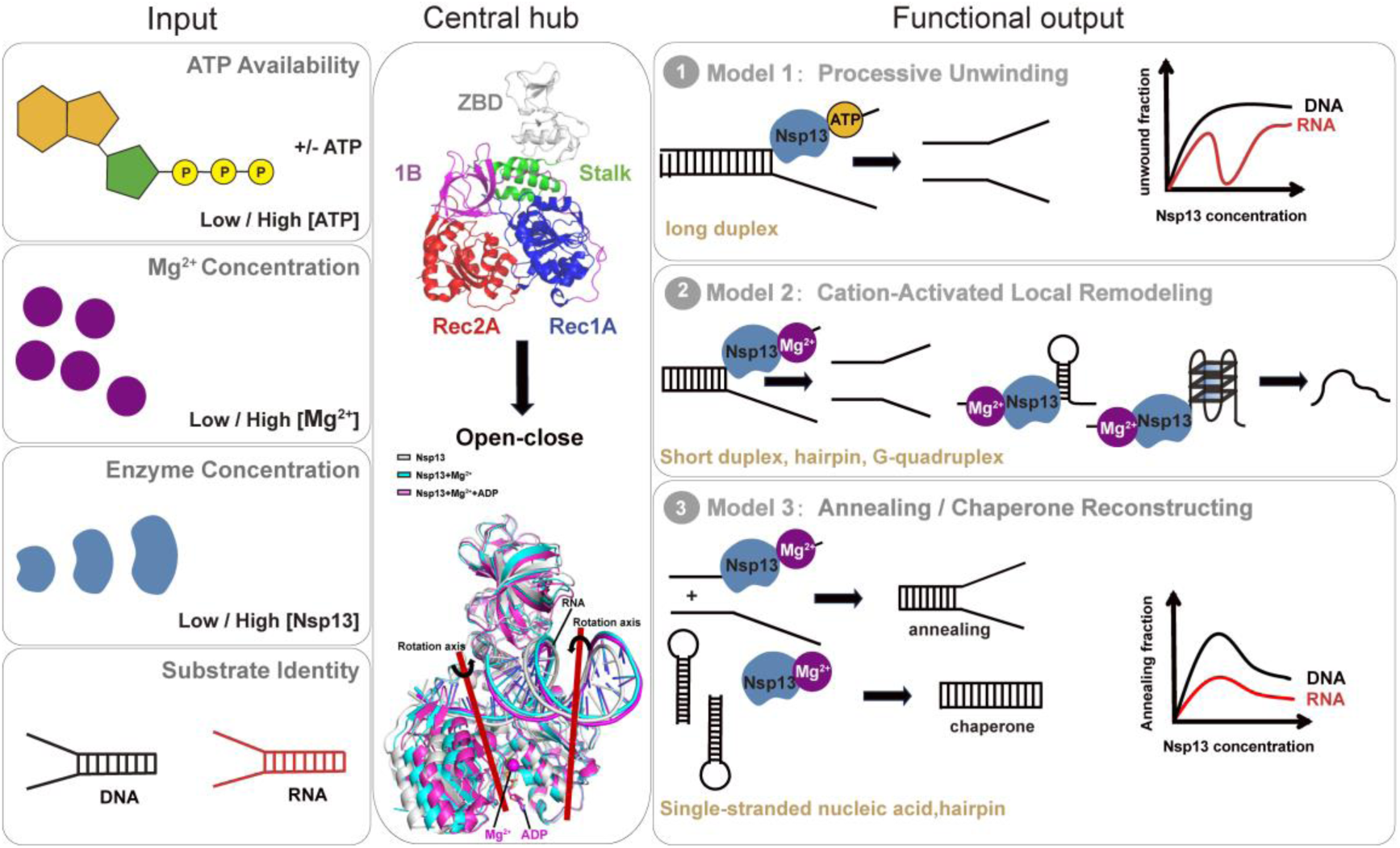
An integrated functional model of Nsp13 as a tunable nucleic acid remodeler. Nsp13 integrates key inputs—cofactors (ATP/Mg²⁺), enzyme concentration, and substrate topology—through a central regulatory hub where Mg²⁺ or Mg²⁺-nucleotide complex allosterically stabilizes its compact RecA1–RecA2 conformation. This tunable hub drives three distinct functional outputs: (1) ATP-dependent processive unwinding, (2) Mg²⁺-primed ATP-independent remodeling, and (3) ATP-independent strand annealing and chaperoning. The coordinated switching among these outputs enables Nsp13 to operate as a multifunctional, context-sensitive remodeler that supports diverse nucleic-acid transactions during viral replication and transcription.

This integrated framework reconciles and extends prior observations on coronavirus helicases. SARS-CoV Nsp13 exhibits ADP-stimulated chaperone activity[24], and SARS-CoV-2 Nsp13 has been reported to resolve RNA stem-loops via ATP-independent mechanisms[25]. Our data provide a unifying mechanistic basis by showing that Mg²⁺ allosterically stabilizes compact RecA1–RecA2 configurations, enabling ATP-independent remodeling while modulating ATP-dependent motor activity. Nsp13 therefore functions not as a single-mode enzyme but as an integrated system whose catalytic output emerges from dynamic conformational regulation.

A key advance of this work is the identification of a robust ATP-independent remodeling mode activated by divalent cations. This activity does not result from nonspecific substrate destabilization, as Mg²⁺ alone stabilizes duplex DNA. Instead, structural modeling and thermal analyses indicate that Mg²⁺ acts as an allosteric effector that primes Nsp13 for low-processivity strand displacement. Preservation of this activity in ATPase-deficient mutants and its enhancement by non-hydrolyzable nucleotides support a hierarchical activation model in which metal-ion binding establishes a remodeling-competent state, while nucleotide binding fine-tunes activity levels. This ATP-independent mode has profound potential implications for the viral lifecycle. It could allow Nsp13 to remain functionally active in cellular niches with low ATP availability or to perform "quick" structural adjustments on nucleic acids without the energetic cost of ATP hydrolysis.

Our findings further demonstrate that unwinding polarity and functional output of Nsp13 are not intrinsic constants but emergent properties shaped by substrate architecture and energetic context. Duplex DNA supports bidirectional remodeling via both ATP-dependent and ATP-independent mechanisms, consistent with cryo-EM observations of 3′→5′ RNA unwinding within the replication–transcription complex[26]. In contrast, G4s show a strong 5′→3′ bias, suggesting higher-order structures impose geometric constraints. RNA substrates additionally reveal pronounced Mg²⁺ sensitivity and nonlinear concentration dependence, highlighting the importance of substrate-specific regulatory logic in governing Nsp13 activity.

Together, these properties provide SARS-CoV-2 with a versatile mechanism to manage its large and structurally complex RNA genome. By integrating multiple remodeling activities, Nsp13 can both resolve obstructive secondary structures and G4s ahead of the replication fork and rescue misfolded RNA intermediates. Notably, the strand-annealing activity may play a key role during discontinuous transcription[37], potentially facilitating template switching essential for subgenomic RNA synthesis[38]. By consolidating these functions within a single protein, Nsp13 reduces reliance on multiple specialized host factors, thereby enhancing viral replication efficiency and autonomy. More broadly, our findings reveal that helicases can encode far greater functional versatility than implied by classical motor-centric definitions, highlighting new conceptual and therapeutic opportunities for targeting viral replication through noncanonical helicase mechanisms.

## Materials and Methods

### Nucleic acid substrates

DNA oligonucleotides were synthesized and HPLC-purified by Sangon Biotech; RNA oligonucleotides by Jinsirui Biotechnology. Sequences and modifications are listed in Table S1. DNA and RNA duplexes were prepared by annealing a FAM-labeled strand (2 µM) with 1.2-fold excess of complementary strand in 20 mM Tris-HCl pH 7.5, 50 mM NaCl, heating at 95°C for 10 min, and cooling to room temperature overnight. G4s were folded by heating 2 µM DNA or RNA in 20 mM Tris-HCl pH 7.5, 100 mM KCl, followed by slow cooling. Hairpins were heat-denatured at 95°C for 5 min and snap-cooled on ice.

### Protein expression and purification

SARS-CoV-2 Nsp13 (GenBank: OR099200.1) was cloned into pET28a with an N-terminal His₆ tag. Plasmids were transformed into *E. coli* Rosetta (DE3) and grown at 37°C on a shaking incubator at 200 rpm for 3 hours until OD₆₀₀ reached approximately 0.8. At this point, cultures were rapidly cooled in an ice-water bath and then incubated at 18°C for 20 hours for self-induction. Cells were lysed in 25 mM HEPES pH 7.0, 500 mM NaCl, 20% glycerol, 5 mM imidazole, 2 mM EDTA, 2 mM DTT. Lysates were applied to His-tag resin, washed with 800 mM NaCl, eluted with 250 mM imidazole, and further purified on a HiTrap Heparin column. Proteins were concentrated, aliquoted, and stored at −80°C. ATPase-deficient mutant K288R and a His-tag-free Nsp13 variant (generated by inserting a SUMO protease cleavage site between the His-tag and Nsp13) were purified using the same protocol.

### LC-MS/MS analysis

Purified Nsp13 was digested with trypsin at 37°C for 12 h. Peptides were desalted, dissolved in 0.1% formic acid, and analyzed on a Fusion Lumos mass spectrometer with PepMap C18 trap (300 μm × 35 mm) and analytical column (75 μm × 150 mm). Data were processed with Proteome Discoverer 2.2 (FDR: 1% strict, 5% relaxed).

### Nucleic acid binding assay

Fluorescence anisotropy was measured on an Infinite F200 (Tecan)[27]. Increasing concentrations of Nsp13 were titrated into 5 nM FAM-labeled nucleic acid in 25 mM Tris-HCl pH 7.5, 20 mM NaCl or KCl (for G4s), 2 mM DTT, and incubated at 37°C for 5 min. Data were fitted to the Hill equation to determine Kd and Hill coefficient (n).

### ATPase activity assay

ATP hydrolysis was measured using the Malachite Green Phosphate Assay Kit (Sigma). Reactions (40 µL) contained 25 mM Tris-HCl pH 7.5, 20 mM NaCl, 5 mM MgCl₂, 2 mM DTT, 1 µM nucleic acid, 1 mM ATP, 400 nM Nsp13, incubated at 37°C for 5 min. Pi release was quantified at 620 nm.

### Helicase unwinding assay

Unwinding reactions were performed by incubating Nsp13 with 40 nM dsDNA or dsRNA substrate in unwinding buffer (25 mM Tris-HCl pH 7.5, 20 mM NaCl, 2 mM DTT) at 37°C for 30 min. Reactions included 0–10 mM MgCl₂, 0–10 mM ATP, and a 20-fold molar excess of unlabeled trap strand (Table S1). DNA unwinding was stopped with 10 × stop buffer (50 mM Tris pH 7.5, 50% glycerol, 1% SDS, 0.1% bromophenol blue). RNA reactions were terminated with 10 × stop buffer (50 mM Tris pH 7.5, 50% glycerol, 2% SDS, 100 mM EDTA, 0.1% bromophenol blue) and digested with 20 mg/mL proteinase K. Products were resolved on 10% (DNA) or 15% (RNA) native PAGE gels, visualized with a ChemiDoc MP system (Bio-Rad), and quantified using ImageJ.

### G4 unfolding assay

DG4 (25-30 nM) or RG-1 (60 nM) was incubated with Nsp13 in substrate-specific buffers (DG4: 25 mM Tris-HCl pH 7.5, 20 mM KCl, 5 mM Mg²⁺; RG-1: 25 mM Tris-HCl pH 7.5, 20 mM KCl, 1 mM Mg²⁺). Reactions were initiated with 1 mM ATP and 30–36 nM trap strand, incubated for 3 min, stopped with 10 × stop buffer (50 mM Tris pH 7.5, 50% glycerol, 1% SDS, 0.1% bromophenol blue), followed by addition of a 20-fold C-trap.

### Nucleic acid annealing assay

Reactions contained 20 nM substrate and complementary strand in 25 mM Tris-HCl pH 7.5, 20 mM NaCl. Specific conditions included 5 mM Mg²⁺ for Mg²⁺-dependent annealing, 2 mM ATP for ATP-dependent annealing, or 1 mM Mg²⁺ for RNA-specific annealing. Reactions were initiated by Nsp13 addition, stopped with 10 × stop buffer (100 mM EDTA, 2% SDS, 50% glycerol, 0.1% bromophenol blue) and 20-fold trap strand, resolved on 10% (DNA) or 15% (RNA) native PAGE.

### Chaperone activity

Pre-folded hairpin DNA or RNA (20 nM each, one FAM-labeled) was incubated with 3 µM Nsp13 in 25 mM Tris-HCl pH 7.5, 20 mM NaCl, 2 mM DTT, 5 mM MgCl₂ (DNA) or 1 mM MgCl₂ (RNA) at 37°C for 1 h. Reactions were stopped with 10× stop buffer (50 mM Tris pH 7.5, 50% glycerol, 2% SDS, 100 mM EDTA, 0.1% bromophenol blue) and analyzed by native PAGE.

### FRET melting

FAM/HEX-labeled duplex DNA (0.5 µM) was melted from 25°C to 95°C on a Rotor-Gene Q (Qiagen). FAM emission was normalized, and Tm was defined at 0.5 normalized emission.

### Circular dichroism (CD) thermal denaturation

Protein stability was measured on a Jasco J-1500 spectropolarimeter. Nsp13 (3 µM) in unwinding buffer was heated from 25°C to 95°C at 2°C/min. CD spectra were recorded from 200–260 nm (1 nm bandwidth); melting curves at 222 nm were fitted to the Boltzmann equation to determine Tm.

### Alphafold3 prediction

Structures of SARS-CoV-2 Nsp13 and its K288R variant were predicted using the AlphaFold3 server. Select predictions included Mg²⁺, ATP (or analogs), and RNA duplexes or hairpins as ligands. Only predictions meeting the following quality criteria were analyzed: pLDDT > 90 for all ligand-coordinating residues, pTM ≥ 0.9, and ipTM ≥ 0.7.

## Author contributions

Xi-Miao Hou: Conceptualization, Supervision, Writing- Reviewing and Editing. Hai-Hong Li: Investigation, Data curation, Writing- Original draft preparation. Jia-Li Hou, Xue-Yang Yu, Jie Jin: Investigation.

## Data availability

All data presented in this study are available upon request.

---

**Table S1.**
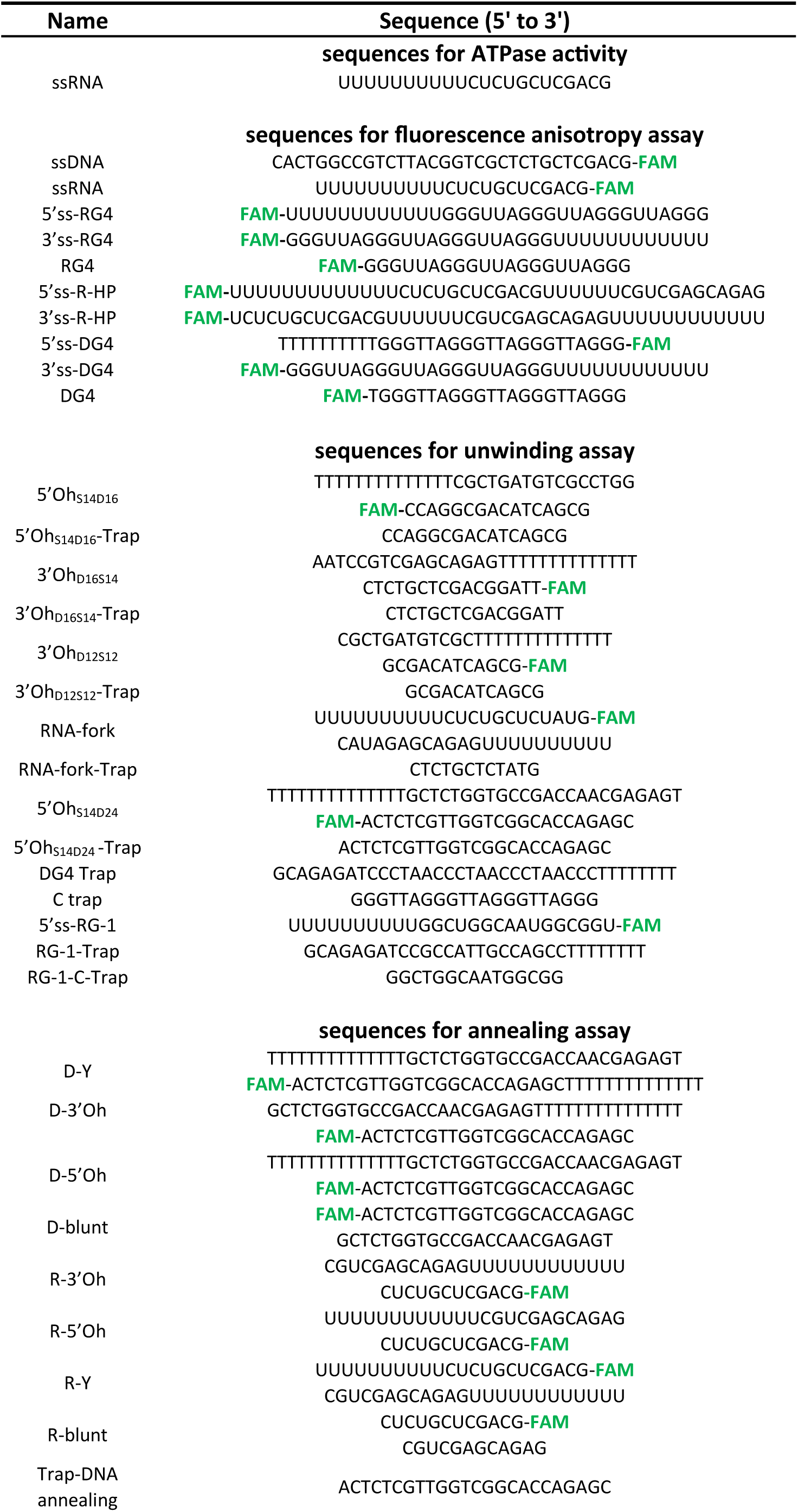
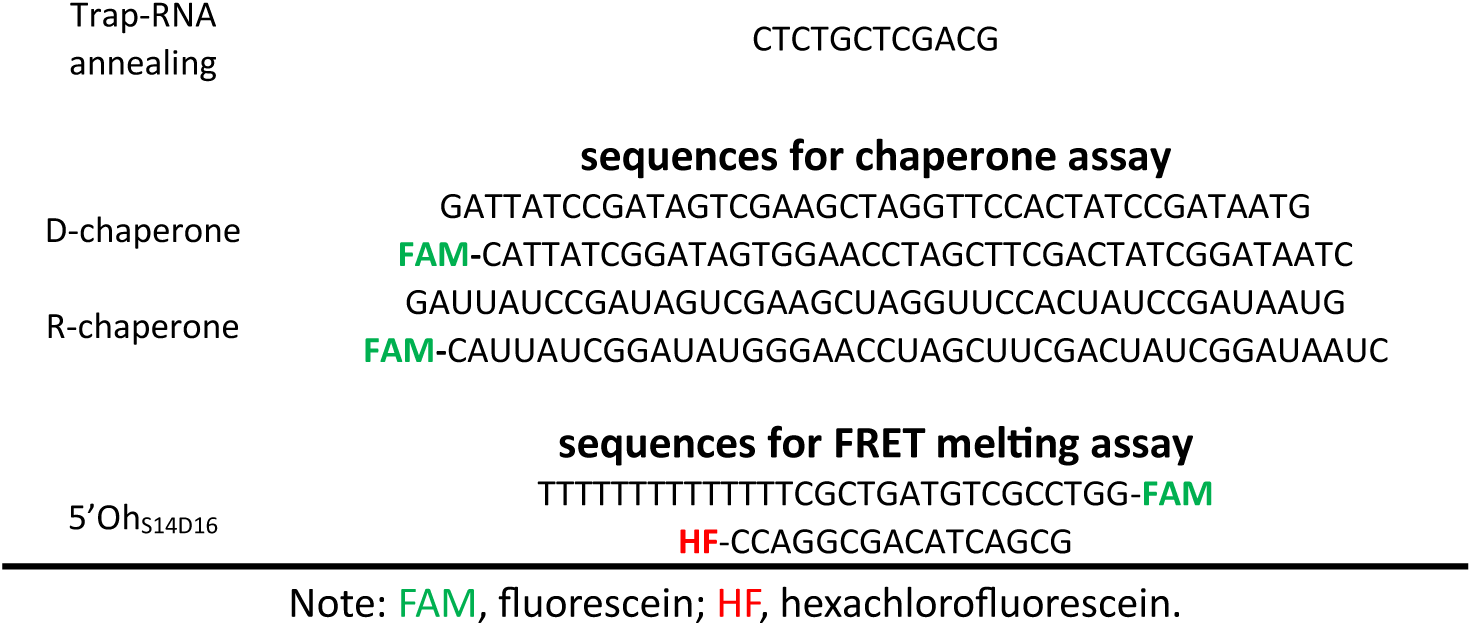
DNA and RNA sequences used in different assays.

**Table S2.**
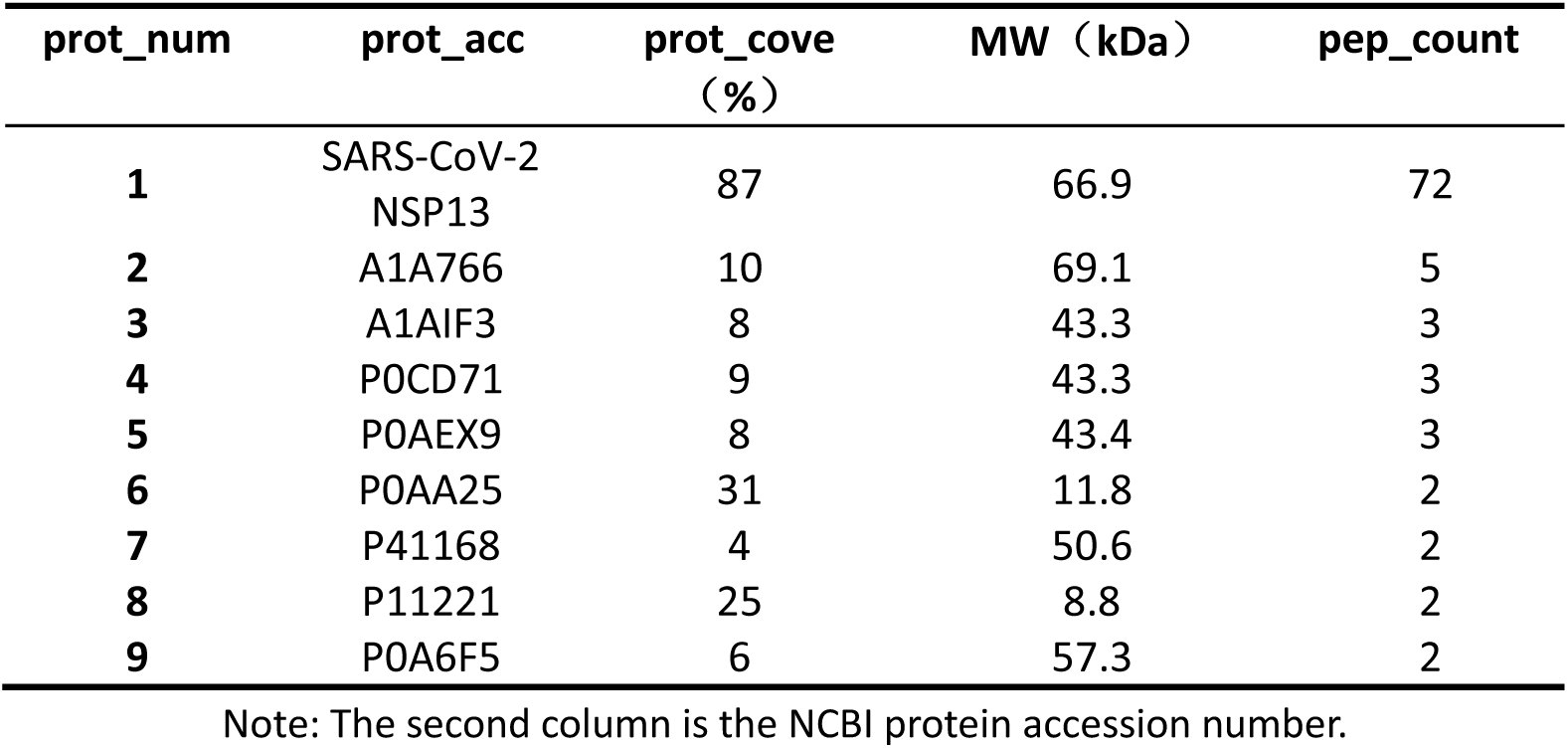
SARS-CoV-2 Nsp13 mass spectrometry.

**Table S3.** Binding parameter of SARS-CoV-2 Nsp13.

| Substrate | Binding parameters |  |
| --- | --- | --- |
|  | *Kd*(nM) | *n* |
| ssDNA | 15.83±1.26 | 1.3 |
| ssRNA | 19.85±0.90 | 2.0 |
| 5'ss-RG4 | 11.60±0.79 | 1.6 |
| 3'ss-RG4 | 13.1±0.81 | 2.3 |
| RG4 | 20.4±3.94 | 0.9 |
| 5'ss-R-HP | 10.85±0.43 | 2.7 |
| 3'ss-R-HP | 33.02±0.40 | 1.1 |
| 5'ss-DG4 | 21.3±0.70 | 1.3 |
| 3'ss-DG4 | 26.0±0.58 | 2.3 |
| DG4 | 51.6±3.56 | 1.6 |

**Figure S1.**
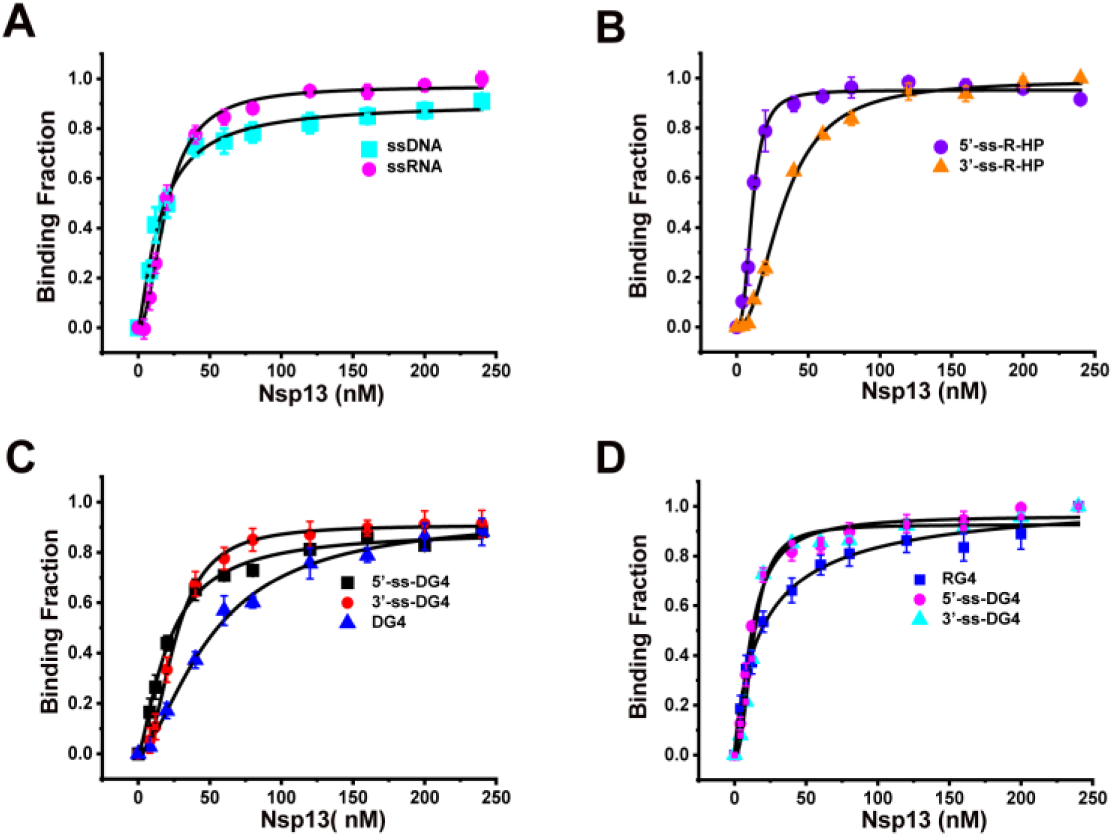
Binding affinity of Nsp13 to various nucleic acid substrates measured by fluorescence anisotropy. Binding isotherms of Nsp13 to (A) single-stranded DNA (ssDNA) and single-stranded RNA (ssRNA); (B) RNA hairpin; (C) DNA G-quadruplex (DG4); and (D) RNA G-quadruplex (RG4). Data are presented as mean ± SD from three independent experiments. Curves were fitted to the Hill equation to determine the equilibrium dissociation constant (*K*d) and Hill coefficient (n).

**Figure S2.**
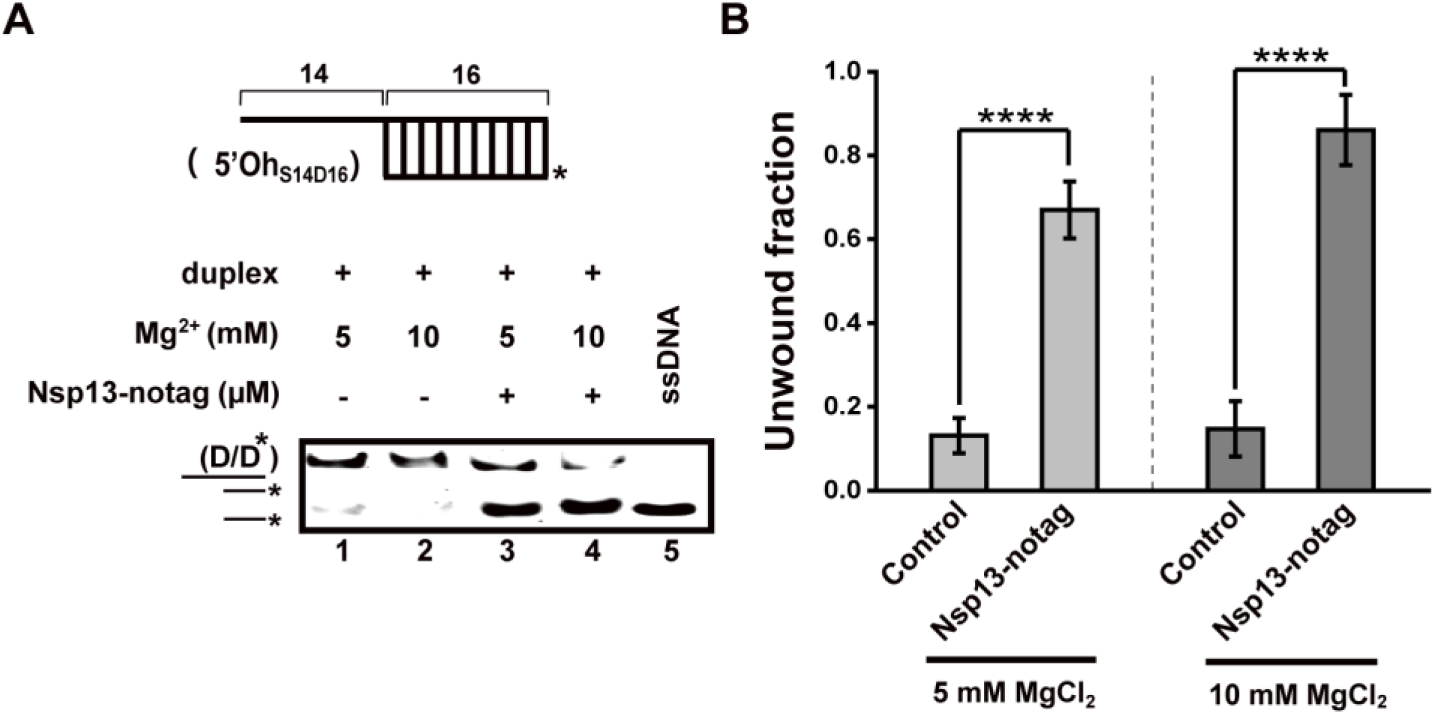
Verification of ATP-independent DNA unwinding by His-tag-free Nsp13. (A) EMSA showing ATP-independent DNA unwinding by His-tag-free Nsp13 (Nsp13-notag). (B) Quantification of unwound DNA fraction illustrating the effect of Mg²⁺ concentration on ATP-independent unwinding. Data are presented as mean ± SD (n=3). Statistical significance was determined by unpaired Student’s t-test.

**Figure S3.**
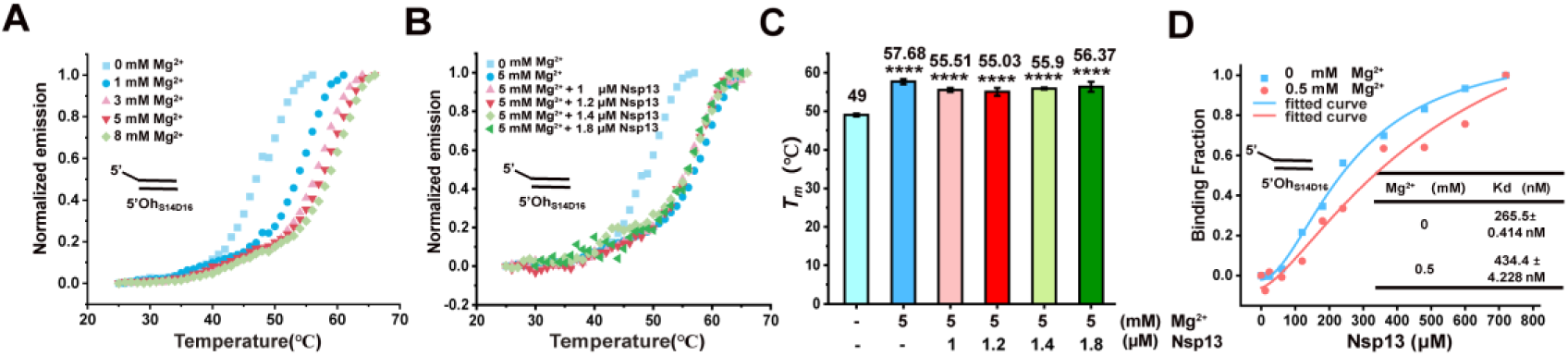
Mechanism of Mg²⁺-mediated DNA unwinding by Nsp13. (A)FRET-melting curves of duplex DNA in the presence of increasing Mg²⁺ concentrations (0–8 mM). (B) FRET-melting curves of duplex DNA in the presence of increasing Nsp13 concentrations (0–1.8 µM). (C) Bar graph showing that Nsp13 does not destabilize the duplex DNA substrate. (D) Binding curves of Nsp13 to duplex DNA in the presence of 0 mM and 0.5 mM Mg²⁺.

**Figure S4.**
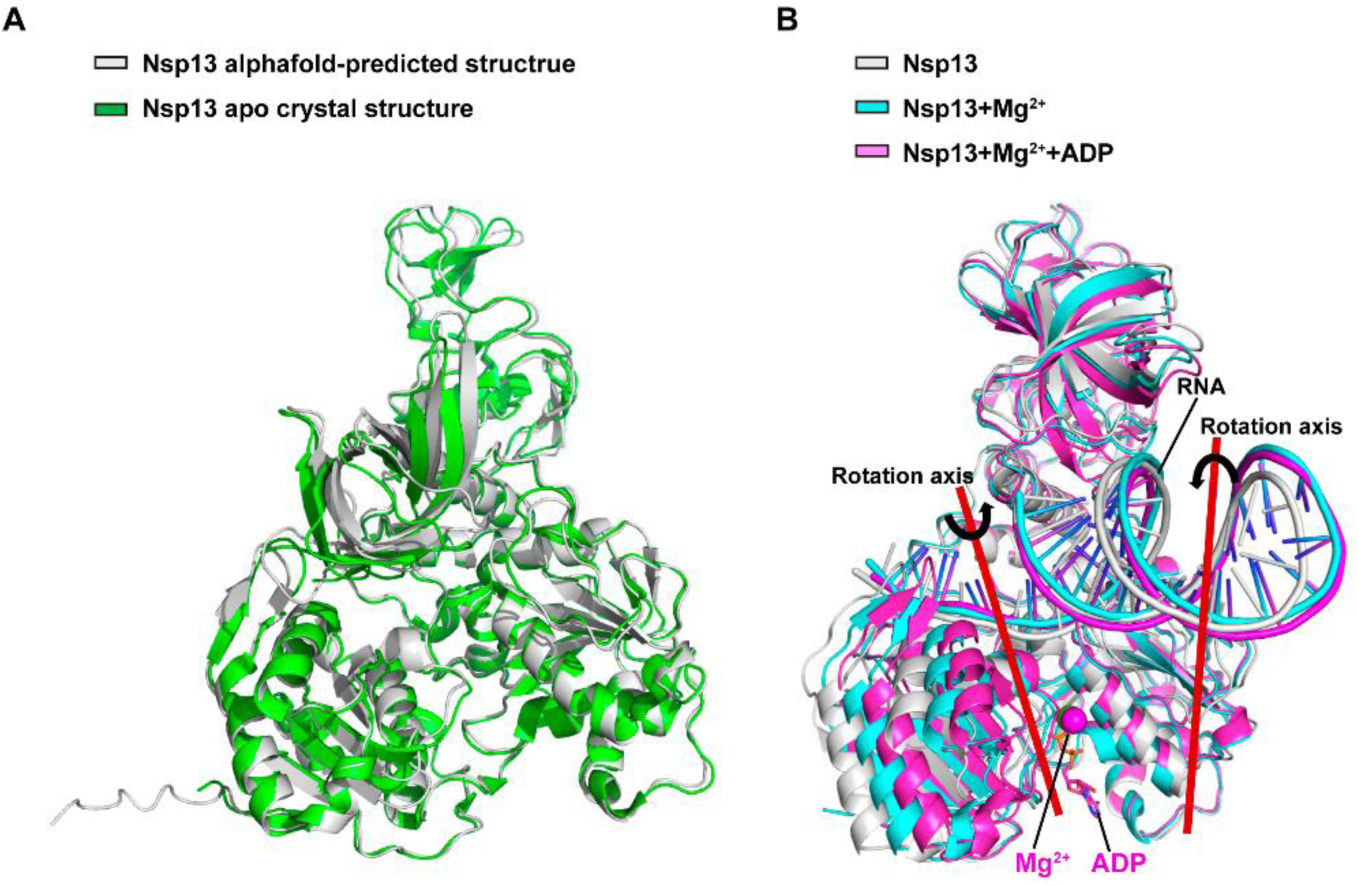
AlphaFold3-predicted Nsp13. (A) The predicted structure of apo Nsp13 aligns well with the experimental structure 7nio from the PDB. (B) Nsp13-ss/dsRNA complex with Mg²⁺ and ADP. Mg²⁺ binds in the RecA1–RecA2 cleft and induces structural compaction. ADP binding further stabilizes the closed conformation.

**Figure S5.**
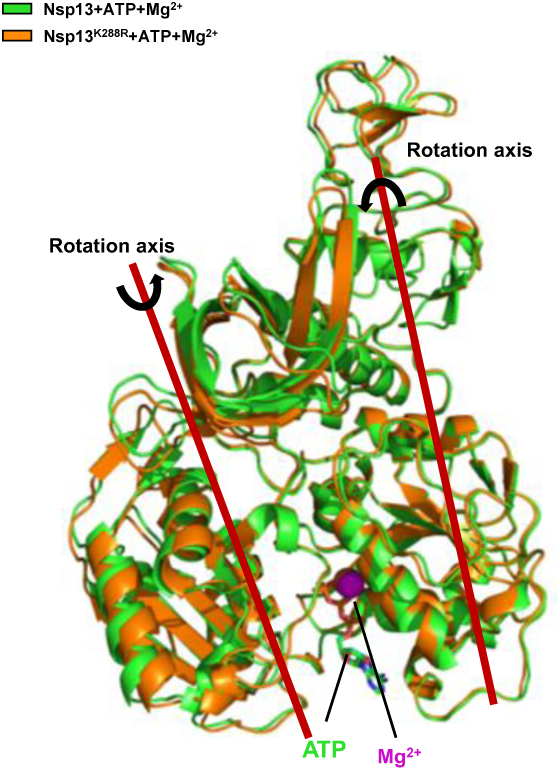
AlphaFold3-predicted structures of Nsp13-ATP-Mg²⁺ and K288R-ATP-Mg²⁺ complexes. Cartoon representation: Nsp13-ATP-Mg²⁺ in green, K288R-ATP-Mg²⁺ in orange. ATP molecules are shown as sticks, and Mg²⁺ ions as purple spheres.

**Figure S6.**
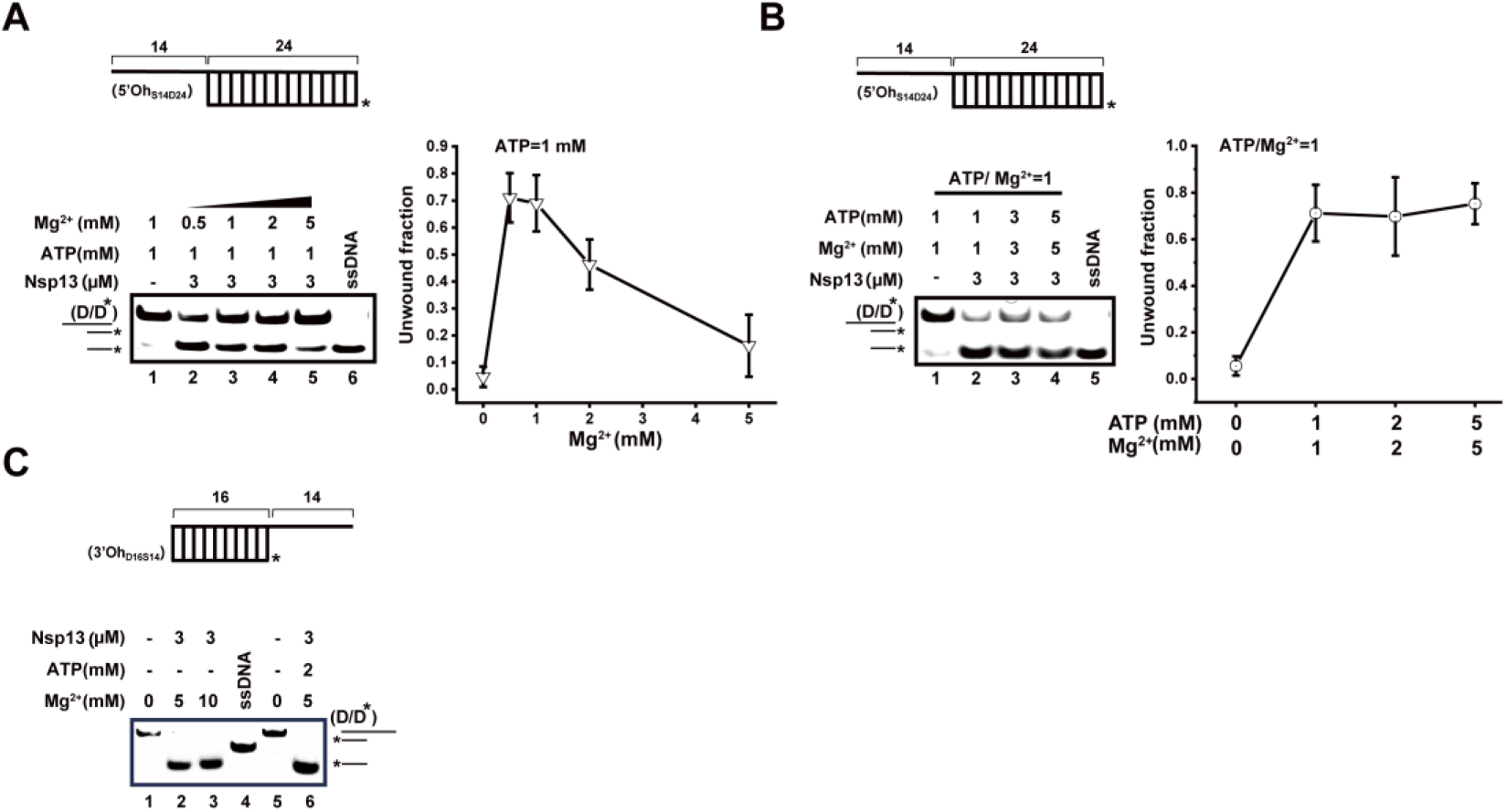
The ATP/Mg²⁺ ratio fine-tunes Nsp13 unwinding of duplex DNA and Nsp13 exhibits 3′→5′ unwinding activity. (A) EMSA (left) and quantification (right) demonstrating that increasing Mg²⁺ concentrations inhibit Nsp13-mediated unwinding at a fixed ATP level (1 mM). The 5′-overhang substrate 5′Oh_S14D24_ (40 nM) was used. (B) Balanced ATP/Mg²⁺ sustains unwinding efficiency. EMSA (left) and quantification (right) showing that maintaining a 1:1 stoichiometry between ATP and Mg²⁺ preserves robust unwinding across a range of ATP concentrations (1–5 mM). All data represent mean ± SD from three independent experiments. (C) Schematic of the 3′-overhang DNA substrate (3′Oh_D16S14_). EMSA illustrates efficient unwinding by Nsp13 in the presence of 5–10 mM Mg²⁺, either without or with 2 mM ATP. Note: The FAM-labeled ssDNA strand migrates more slowly than the enzyme-released unwound product, likely due to secondary structure formation in the absence of Nsp13.

**Figure S7.**
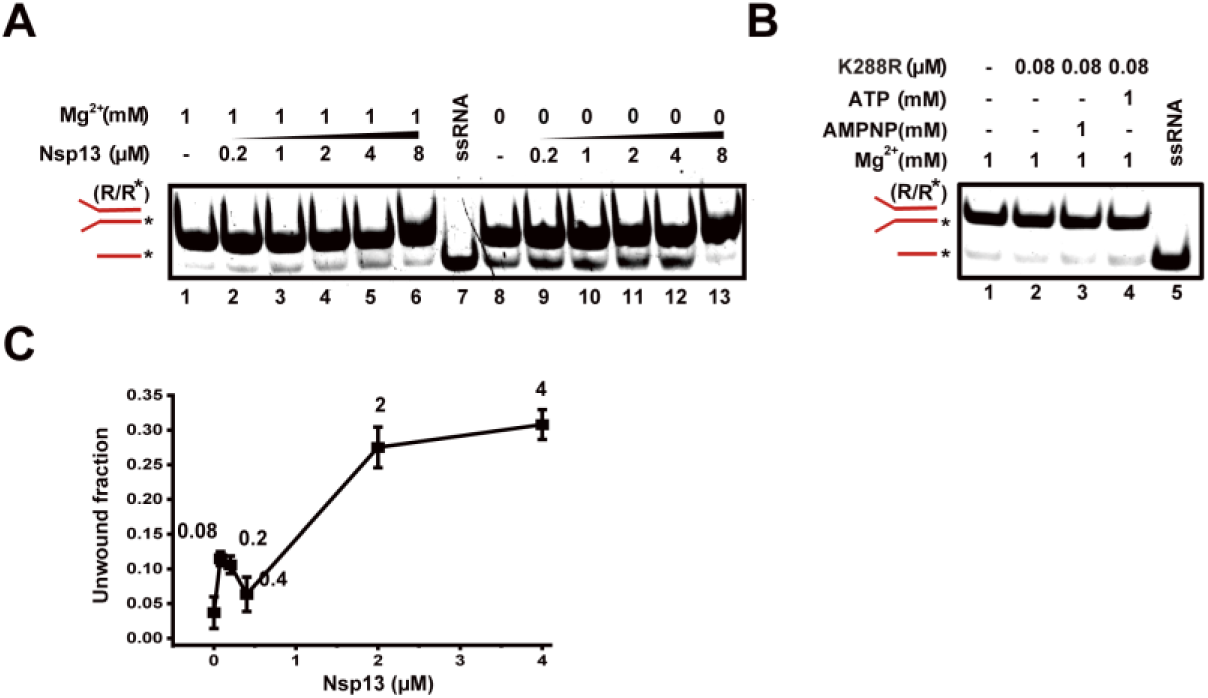
Mg²⁺ and Nsp13 concentration dependence of RNA fork unwinding. (A) EMSA showing RNA unwinding by Nsp13 in the absence of ATP at 0 mM and 1 mM Mg²⁺. (B) EMSA analysis of RNA unwinding by the ATPase-deficient mutant K288R at low Nsp13 concentration (80 nM) under different ATP conditions. (C) Quantification of unwound RNA fractions at 0.5 mM Mg²⁺ as a function of Nsp13 concentration, reproduced from Figure 4A. Data represent mean ± SD from three independent replicates.

**Figure S8.**
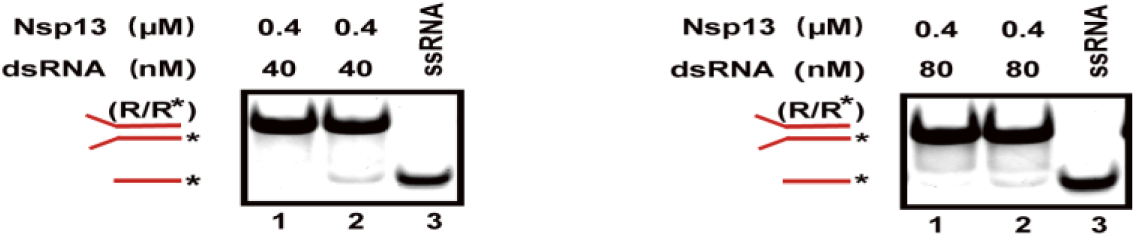
Analysis of RNA duplex unwinding under inhibitory Nsp13 conditions. EMSA showing RNA duplex unwinding at the same Nsp13 concentration (0.4 μM) that inhibited RNA unwinding in Figure 4H in the presence of 1 mM ATP–Mg²⁺, but using a different RNA substrate concentration (40 nM vs. 80 nM).

**Figure S9.**
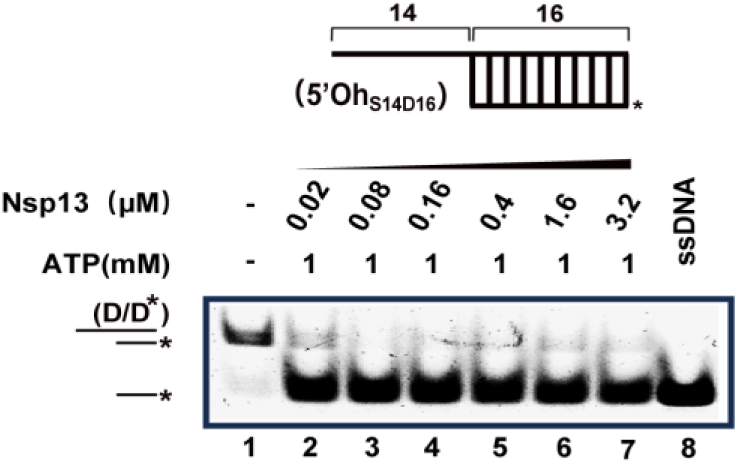
ATP-dependent DNA unwinding by Nsp13 increases with protein concentration. (A) EMSA analysis of helicase activity on duplex DNA with increasing Nsp13 concentrations (0.02–3.2 µM). Reaction conditions: 40 nM DNA, 1 mM Mg^2+^, 1 mM ATP, 37°C. (B) Quantification of unwound DNA fractions. Data represent mean ± SD from three replicates.

**Figure S10.**
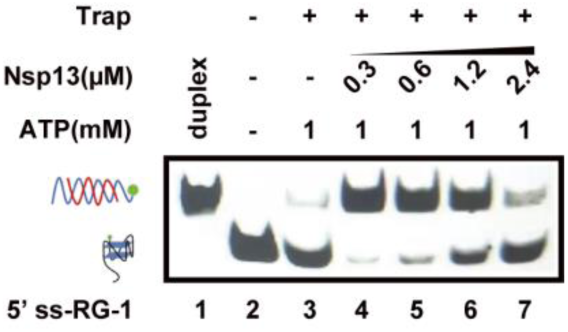
EMSA analysis of SARS-CoV-2 RNA G-quadruplex (RG-1) unfolding by varying concentrations of Nsp13 in 1 mM Mg²⁺. Partial unfolding was observed at low to moderate protein concentrations, whereas higher Nsp13 levels inhibited G4 unfolding.

**Figure S11.**
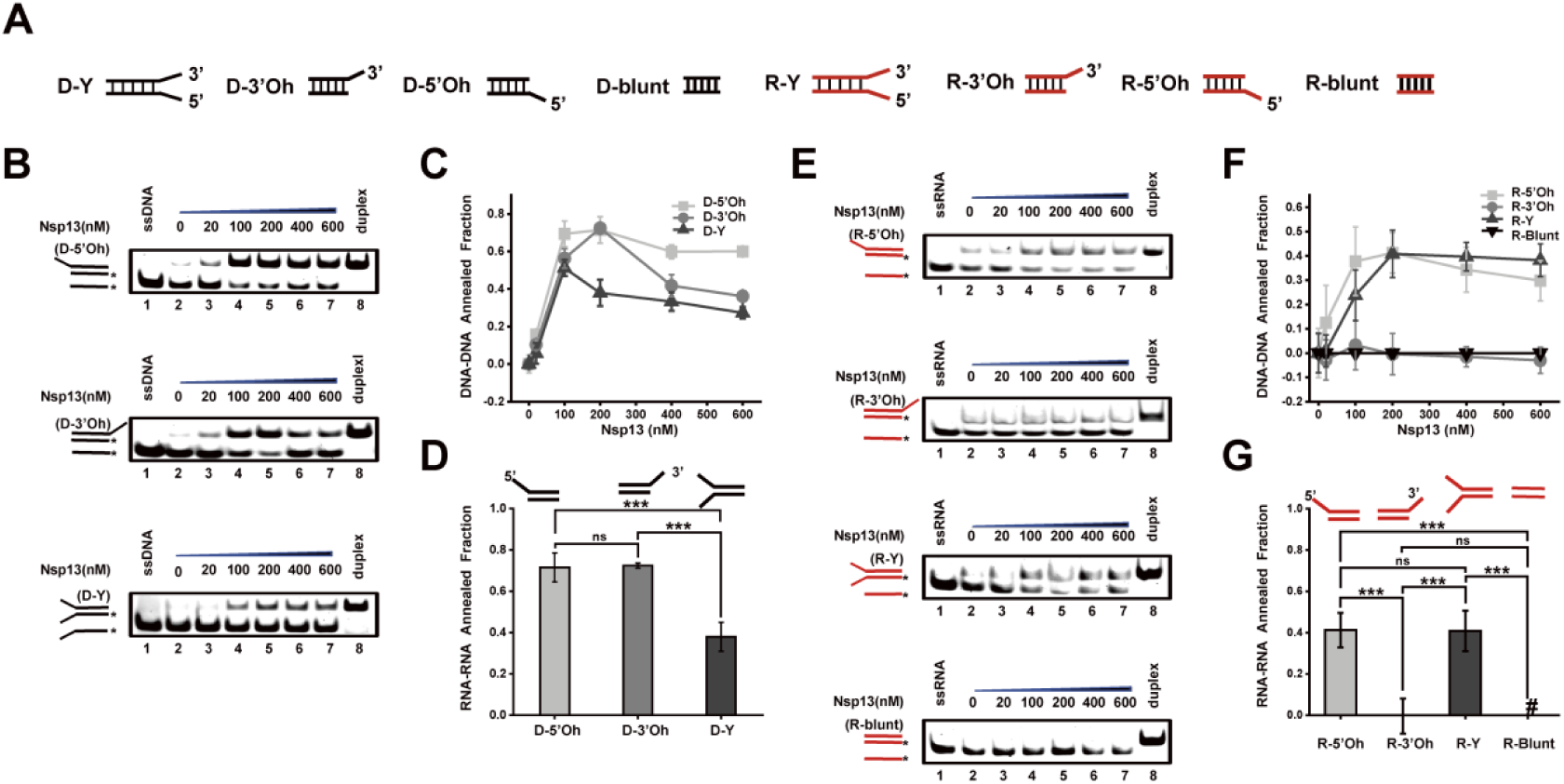
Nsp13 exhibits ATP- and Mg²⁺-independent strand annealing activity on diverse DNA and RNA substrates. (A) Schematic of DNA and RNA substrate structures used in annealing assays. (B) EMSA results showing Nsp13-mediated annealing of DNA substrates with 5′-overhang, 3′-overhang, and fork structures, respectively. (C) Concentration–response curve of DNA annealing activity in the absence of Mg^2+^ and ATP. (D) Annealing efficiency for three DNA configurations at 200 nM Nsp13. (E) EMSA results showing RNA annealing with 5′-overhang, 3′-overhang, fork, and blunt-end substrates. (F) Concentration–response curve of RNA annealing activity in the absence of Mg^2+^ and ATP. (G) Annealing efficiency for four RNA configurations at 200 nM Nsp13. Data are presented as mean ± SD (n=3). Statistical significance was determined by one-way ANOVA with Tukey’s HSD test; **^***^**P < 0.001; ns, not significant. Note: DNA blunt-end substrates were excluded from the analysis due to instability under the annealing buffer conditions.

**Figure S12.**
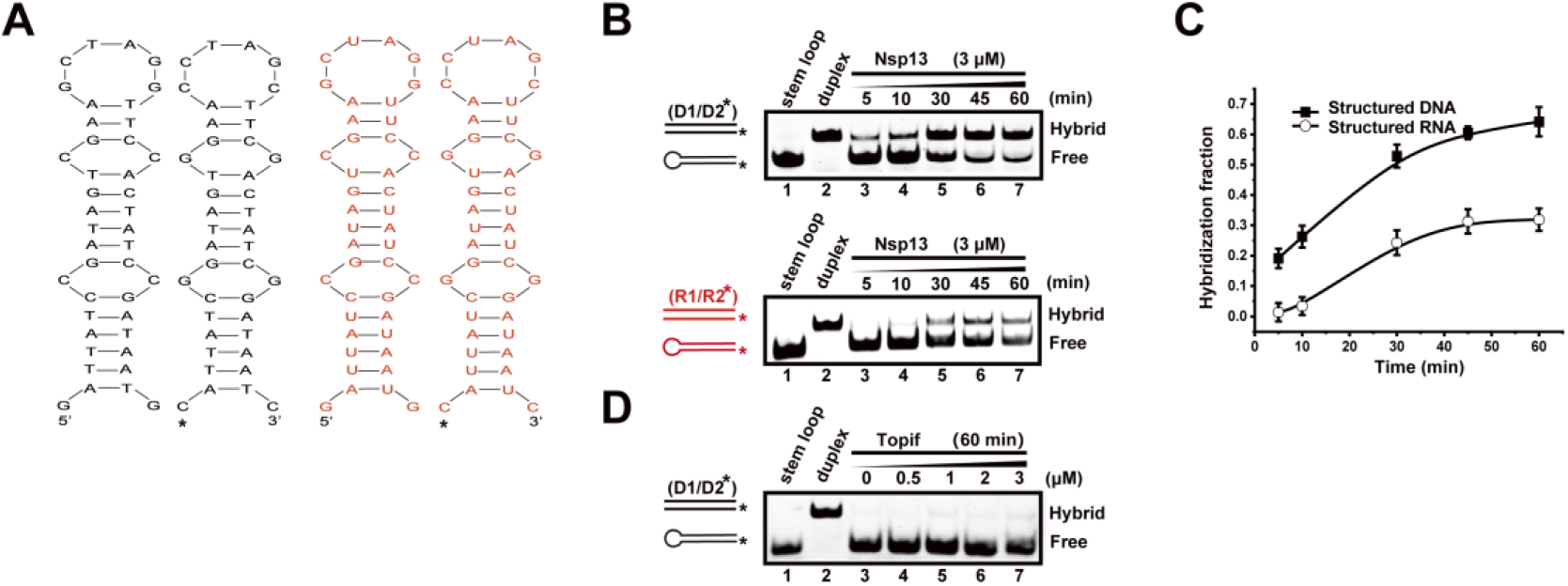
Chaperone activity of Nsp13. (A) Schematic of the strand-exchange assay. Complementary 42-nt DNA and RNA stem-loops were used as substrates, with the FAM-labeled strands indicated by asterisks. (B–C) Time-course EMSA (B) and quantification (C) showing Nsp13-mediated strand exchange in the absence of ATP, demonstrating its intrinsic chaperone activity. Hybridization fraction was calculated from three independent replicates (mean ± SD). (D) Control experiment with ToPif1, a canonical 5′→3′ helicase. Pre-folded DNA stem-loops were incubated with increasing concentrations of ToPif1 (0–3 µM) under identical conditions. No strand exchange was observed, confirming that the chaperone activity is unique to Nsp13 and not a general property of helicases. All reactions were performed with 20 nM FAM-labeled and unlabeled strands.

## References

[1] . Kim, D., et al., The Architecture of SARS-CoV-2 Transcriptome. Cell, 2020. 181(4): p. 914–921.e10.

[2] . Huston, N.C., et al., Comprehensive in vivo secondary structure of the SARS-CoV-2 genome reveals novel regulatory motifs and mechanisms. Molecular cell, 2021. 81(3): p. 584–598.e5.

[3] . Ji, D., et al., Discovery of G-quadruplex-forming sequences in SARS-CoV-2. Briefings in bioinformatics, 2021. 22(2): p. 1150–1160.

[4] . Zhai, L., et al., Recent advances in applying G-quadruplex for SARS-CoV-2 targeting and diagnosis: A review. International journal of biological macromolecules, 2022. 221: p. 1476–1490.

[5] . Zhai, L., et al., Targeting the RNA G-Quadruplex and Protein Interactome for Antiviral Therapy. Journal of medicinal chemistry, 2022. 65(15): p. 10161–10182.

[6] . Zhao, C., et al., Targeting RNA G-Quadruplex in SARS-CoV-2: A Promising Therapeutic Target for COVID-19? Angewandte Chemie (International ed. in English), 2021. 60(1): p. 432–438.

[7] . Chauhan, S. and S.A. Woodson, Tertiary interactions determine the accuracy of RNA folding. Journal of the American Chemical Society, 2008. 130(4): p. 1296–1303.

[8] . Musier-Forsyth, K., RNA remodeling by chaperones and helicases. RNA biology, 2010. 7(6): p. 632–633.

[9] . Jarmoskaite, I. and R. Russell, RNA helicase proteins as chaperones and remodelers. Annual review of biochemistry, 2014. 83: p. 697–725.

[10] . Yang, J., et al., RNA chaperones encoded by RNA viruses. Virologica Sinica, 2015. 30(6): p. 401–409.

[11] . Bai, C., Q. Zhong and G.F. Gao, Overview of SARS-CoV-2 genome-encoded proteins. Science China. Life sciences, 2022. 65(2): p. 280–294.

[12] . Chen, J., et al., Structural Basis for Helicase-Polymerase Coupling in the SARS-CoV-2 Replication-Transcription Complex. Cell, 2020. 182(6): p. 1560–1573.e13.

[13] . Shu, T., et al., SARS-Coronavirus-2 Nsp13 Possesses NTPase and RNA Helicase Activities That Can Be Inhibited by Bismuth Salts. Virologica Sinica, 2020. 35(3): p. 321–329.

[14] . Tanner, J.A., et al., The severe acute respiratory syndrome (SARS) coronavirus NTPase/helicase belongs to a distinct class of 5’ to 3’ viral helicases. The Journal of biological chemistry, 2003. 278(41): p. 39578–39582.

[15] . Jia, Z., et al., Delicate structural coordination of the Severe Acute Respiratory Syndrome coronavirus Nsp13 upon ATP hydrolysis. Nucleic acids research, 2019. 47(12): p. 6538–6550.

[16] . Cao, C., et al., Molecular epidemiology analysis of early variants of SARS-CoV-2 reveals the potential impact of mutations P504L and Y541C (NSP13) in the clinical COVID-19 outcomes. Infection, genetics and evolution : journal of molecular epidemiology and evolutionary genetics in infectious diseases, 2021. 92: p. 104831–104831.

[17] . Grimes, S.L. and M.R. Denison, The Coronavirus helicase in replication. Virus research, 2024. 346: p. 199401–199401.

[18] . White, M.A., W. Lin and X. Cheng, Discovery of COVID-19 Inhibitors Targeting the SARS-CoV-2 Nsp13 Helicase. The journal of physical chemistry letters, 2020. 11(21): p. 9144–9151.

[19] . Ugurel, O.M., et al., Evaluation of the potency of FDA-approved drugs on wild type and mutant SARS-CoV-2 helicase (Nsp13). International journal of biological macromolecules, 2020. 163: p. 1687–1696.

[20] . Perez-Lemus, G.R., et al., Toward wide-spectrum antivirals against coronaviruses: Molecular characterization of SARS-CoV-2 NSP13 helicase inhibitors. Science advances, 2022. 8(1): p. eabj4526–eabj4526.

[21] . Marx, S.K., et al., Observing inhibition of the SARS-CoV-2 helicase at single-nucleotide resolution. Nucleic acids research, 2023. 51(17): p. 9266–9278.

[22] . Sommers, J.A., et al., Biochemical analysis of SARS-CoV-2 Nsp13 helicase implicated in COVID-19 and factors that regulate its catalytic functions. The Journal of biological chemistry, 2023. 299(3): p. 102980–102980.

[23] . Park, J., et al., ATPase-dependent duplex nucleic acid unwinding by SARS-CoV-2 nsP13 relies on facile binding and translocation along single-stranded nucleic acid. The Journal of biological chemistry, 2025. 301(7): p. 110373–110373.

[24] . Yu, J., et al., A novel ADP-directed chaperone function facilitates the ATP-driven motor activity of SARS-CoV helicase. Nucleic acids research, 2025. 53(3): p. gkaf034.

[25] . Dumm, A.J., et al., SARS-CoV-2 point mutations are over-represented in terminal loops of RNA stem-loop structures that can be resolved by Nsp13 helicase in a unique manner with respect to nucleotide dependence. Nucleic acids research, 2025. 53(10): p. gkaf447.

[26] . Yan, L., et al., Structural basis for the concurrence of template recycling and RNA capping in SARS-CoV-2. Cell, 2025. 188(25): p. 7194–7205.e10.

[27] . Dou, S. and X.G. Xi, Fluorometric assays for characterizing DNA helicases. Methods (San Diego, Calif.), 2010. 51(3): p. 295–302.

[28] . Newman, J.A., et al., Structure, mechanism and crystallographic fragment screening of the SARS-CoV-2 NSP13 helicase. Nature communications, 2021. 12(1): p. 4848–4848.

[29] . Maio, N., et al., An iron-sulfur cluster in the zinc-binding domain of the SARS-CoV-2 helicase modulates its RNA-binding and -unwinding activities. Proceedings of the National Academy of Sciences of the United States of America, 2023. 120(33): p. e2303860120–e2303860120.

[30] . Jang, K., et al., A high ATP concentration enhances the cooperative translocation of the SARS coronavirus helicase nsP13 in the unwinding of duplex RNA. Scientific reports, 2020. 10(1): p. 4481–4481.

[31] . Liu, G., et al., RNA G-quadruplex in TMPRSS2 reduces SARS-CoV-2 infection. Nature communications, 2022. 13(1): p. 1444–1444.

[32] . Qin, G., et al., RNA G-quadruplex formed in SARS-CoV-2 used for COVID-19 treatment in animal models. Cell discovery, 2022. 8(1): p. 86–86.

[33] . Hou, X., et al., Involvement of G-triplex and G-hairpin in the multi-pathway folding of human telomeric G-quadruplex. Nucleic acids research, 2017. 45(19): p. 11401–11412.

[34] .Miclot, T., et al., Structure and Dynamics of RNA Guanine Quadruplexes in SARS-CoV-2 Genome. Original Strategies against Emerging Viruses. The journal of physical chemistry letters, 2021. 12(42): p. 10277–10283.

[35] . Yang, J., et al., A cypovirus VP5 displays the RNA chaperone-like activity that destabilizes RNA helices and accelerates strand annealing. Nucleic acids research, 2014. 42(4): p. 2538–2554.

[36] . Dai, Y., et al., Structural mechanism underpinning Thermus oshimai Pif1-mediated G-quadruplex unfolding. EMBO reports, 2022. 23(7): p. e53874–e53874.

[37] . Wang, D., et al., The SARS-CoV-2 subgenome landscape and its novel regulatory features. Molecular cell, 2021. 81(10): p. 2135–2147.e5.

[38] . Kim, D., et al., The Architecture of SARS-CoV-2 Transcriptome. Cell, 2020. 181(4): p. 914–921.e10.

